# RNF144A shapes the hierarchy of cytokine signaling to provide protective immunity against influenza

**DOI:** 10.1101/782680

**Authors:** B. Afzali, S. Kim, E. West, E. Nova-Lamperti, N. Cheru, H. Nagashima, B. Yan, T Freiwald, N. Merle, D. Chauss, M. Bijlmakers, G. Weitsman, Z. Yu, D. Jankovic, S. Mitra, A. Villarino, C. Kemper, A. Laurence, M. Kazemian, J.J. O’Shea, S. John

## Abstract

Cytokine-induced signaling pathways are tightly regulated and self-limiting, as their dysregulation causes immune disorders and cancer. The precise mechanisms that fine-tune these responses are incompletely understood. We show that the E3 ubiquitin ligase *RNF144A* is an IL-2/STAT5-induced gene in T cells and critically orchestrates the hierarchy of IL-2R signaling to promote STAT5 activation and limit RAF-ERK-MAPK output from the IL-2R. Mechanistically, RNF144A increased the interaction between IL-2Rβ and STAT5 and polyubiquitinated RAF1, enhancing its proteasomal degradation and preventing the formation of the potent RAF1/BRAF kinase complex. CD8^+^ T cells from *Rnf144a*^*–/–*^ mice had impaired IL-2-induction of effector genes, including *Tnf* and granzymes, and these mice demonstrated increased susceptibility to influenza. Reduced RNF144A expression was associated with more severe influenza in humans and its expression in patients was a biomarker distinguishing moderate from severe disease. These studies reveal a vital physiological role for RNF144A in maintaining the fidelity of IL-2R signaling in CTLs to prevent severe inflammation in response to infection.

**One Sentence Summary:** RNF144A promotes anti-viral immunity by regulating the hierarchy of cytokine signal output

## Introduction

The fidelity and duration of cytokine-induced signal transduction underpins the fate decisions made by immune cells in response to infections or antigenic challenge. Interleukin-2 (IL-2) is a critically important cytokine with a diverse functional repertoire that includes regulation of both adaptive and innate immunity and maintenance of self-tolerance ^1,2^. For these reasons there is considerable interest in the use of IL-2, the use of anti-IL-2 and recently introduced inhibitors of IL-2 signaling for the clinical management of conditions as diverse as auto-immunity and cancer ^3^. IL-2 is particularly important for orchestrating the effector proteome and function of CD8^+^ cytotoxic T cells (CTLs) compared with CD4^+^ T helper cells and CTL are more acutely responsive to low doses of IL-2 compared with CD4^+^ T helper cells ^4-8^. Indeed, mice lacking STAT5 expression have preferential loss of CD8 versus CD4 T cells, as do mice and non-human primates treated with the JAK1/3 inhibitor tofacitinib ^9,10^.

The high affinity IL-2 receptor (IL-2R) comprises IL-2R*β* and IL-2R*γ* heterodimers, involved in both ligand binding and intracellular signal transduction, and a non-signaling IL-2R*α* (CD25) subunit, which confers high-affinity binding of IL-2 to its receptor ^1^. The association of IL-2 with the IL-2R complex activates three main biochemical downstream signaling pathways, JAK-STAT5, RAS-RAF-MAPK and PI3K-AKT ^2^, which together function as an integrated, cross-regulating network of signal relays, to shape the unique transcriptional output for this cytokine. Mass spectrometry studies show that nearly 80% of IL-2R signaling is dependent on the JAK-STAT5 pathway, but the activation of PI(3,4,5)P_3_-AKT pathway in T cells is independent of IL-2-JAK1/3 signaling, requiring instead constitutively active src family kinases, lck and fyn ^11^. However, the molecular details of the regulators and mechanisms that ensure the hierarchy and fidelity of IL-2-induced signal transduction, arguably key nodes for drug targeting, are incompletely understood. Indeed, although sustaining dominant STAT5 signal output from the IL-2R is critical for key immunoregulatory functions, detailed genetic studies in mice have revealed that STAT5-mediated biological functions are only significantly perturbed when a minimum of three out of four alleles of *Stat5* are deleted, implying that there are mechanisms to maintain dominance of the STAT5 signal even when STAT5 protein is limiting ^12^.

Faithful cytokine signal transduction is exquisitely controlled and “fine-tuned” by precise activation/inactivation events through post-translational modifications of proteins, including phosphorylation and ubiquitination. Ubiquitination of proteins is an important post-translational mechanism to regulate signal transduction, remove unwanted proteins or modify their function and/or subcellular localization ^13,14^. Ubiquitination is mediated by the sequential enzymatic actions of E1 ubiquitin (Ub)-activating enzymes (E1s), E2 Ub-conjugating proteins (E2s) and E3 Ub-ligases (E3s). E3s are vital in this process because they determine substrate specificity and catalyze the transfer of Ub to the substrate from the E2 ^15^. The three main families of E3s are the really interesting new gene (RING), the homologous to E6AP carboxy terminus (HECT) and the ring between ring (RBR), distinguished by their unique mechanism of ubiquitin transfer to substrate ^16,17^. The RBR family are RING-HECT hybrids and have the ability to directly transfer Ub to substrates ^18,19^ and can therefore, function with UbcH7, an E2 that can only function with HECT E3s, as well as with the less restrictive E2 UbcH5a ^18,20^. RBR proteins are highly conserved through evolution and members of this family, such as Parkin, HHARI and LUBAC are associated with neurological disorders, malignant transformation and/or autoimmunity, respectively ^21-24^. RNF144A is a member of the RBR family that has recently been shown to assemble ubiquitin linkages at the K6-, K11- and K48-positions of ubiquitin *in vitro* ^25^, and whose ubiquitin ligase activity is regulated by self-association through its transmembrane domain ^26^. RNF144A plays a role in DNA repair in cancer by targeting DNA-dependent protein kinase (DNA-PK) ^27^ and PARP1 ^28^ for degradation. However, its biological roles in immunity are unknown.

Here we sought to identify and characterize novel regulators of IL-2-induced JAK/STAT5 signaling in immune cells. Through transcriptome and ChIP-seq studies, we identified RNF144A as one of the most highly induced genes regulated by IL-2-STAT5 in primary T cells. Mice with deletion of *Rnf144a* were runted in size and CD8^+^ T cell immunodeficient, exhibiting diminished CTL function, leading to widespread inflammation in response to influenza infection. CTLs from these mice had dysregulated signal output from the IL-2R, showing diminished STAT5 and elevated ERK phosphorylation. Mechanistically, RNF144A was located primarily at the plasma membrane and associated with IL-2Rβ, STAT5 and RAF1. Following IL-2 stimulation, RNF144A enhanced the association between STAT5 and IL-2Rβ, promoting STAT5 activation. Conversely, RNF144A directly targeted RAF1 for proteasomal degradation to limit IL-2-induced RAF-ERK1/2 activation. In human patients, RNF144A expression correlated inversely with severity of influenza infection, being lower in subjects with more severe disease, and with ERK target genes. RNF144A mRNA expression performed well as a biomarker distinguishing severe from moderate cases of influenza. Collectively, our data reveal that the fidelity of cellular IL-2 responsiveness and subsequent development of CTL effector function requires RNF144A-dependent restriction of RAF, and promotion of JAK-STAT5, activation. Thus, RNF144A is essential for fine-tuning IL-2R signaling to shape protective anti-viral responses.

## Results

### IL-2 regulates the T cell ubiquitinome, with RNF144A being the most highly induced IL-2/STAT5-dependent E3 ligase

We used an unbiased transcriptomic approach in pre-activated human CD8^+^ cytotoxic T lymphocytes (CTLs), in the presence and absence of IL-2, with and without a JAK inhibitor (JAKi), to identify key and novel regulators of IL-2-JAK/STAT5 signaling. Approximately 1900 genes were differentially expressed in CTLs treated with IL-2 compared to untreated controls (fold change at least 2, p<0.05) (**Fig. 1a and Fig. S1a, Table S1**). Genes were assigned to functionally grouped terms in ClueGO and ranked by group p-value (**Fig. 1b and Table S2**). The most highly enriched functional category of genes conformed to cell cycle processes and DNA replication. The next highly enriched category of genes controlled by IL-2 was regulation of ubiquitin ligases (**Fig. 1b**). Although ubiquitination is a known mechanism for regulating IL-2R biogenesis and function ^29,30^, the high incidence of genes associated with ubiquitination, approximately three times higher than expected (p = 3.4 × 10^−17^, **Table S2**) in the list of genes regulated by IL-2, was unexpected. Thus, ubiquitinylation is a key cellular process regulated by IL-2.

**Figure 1.**
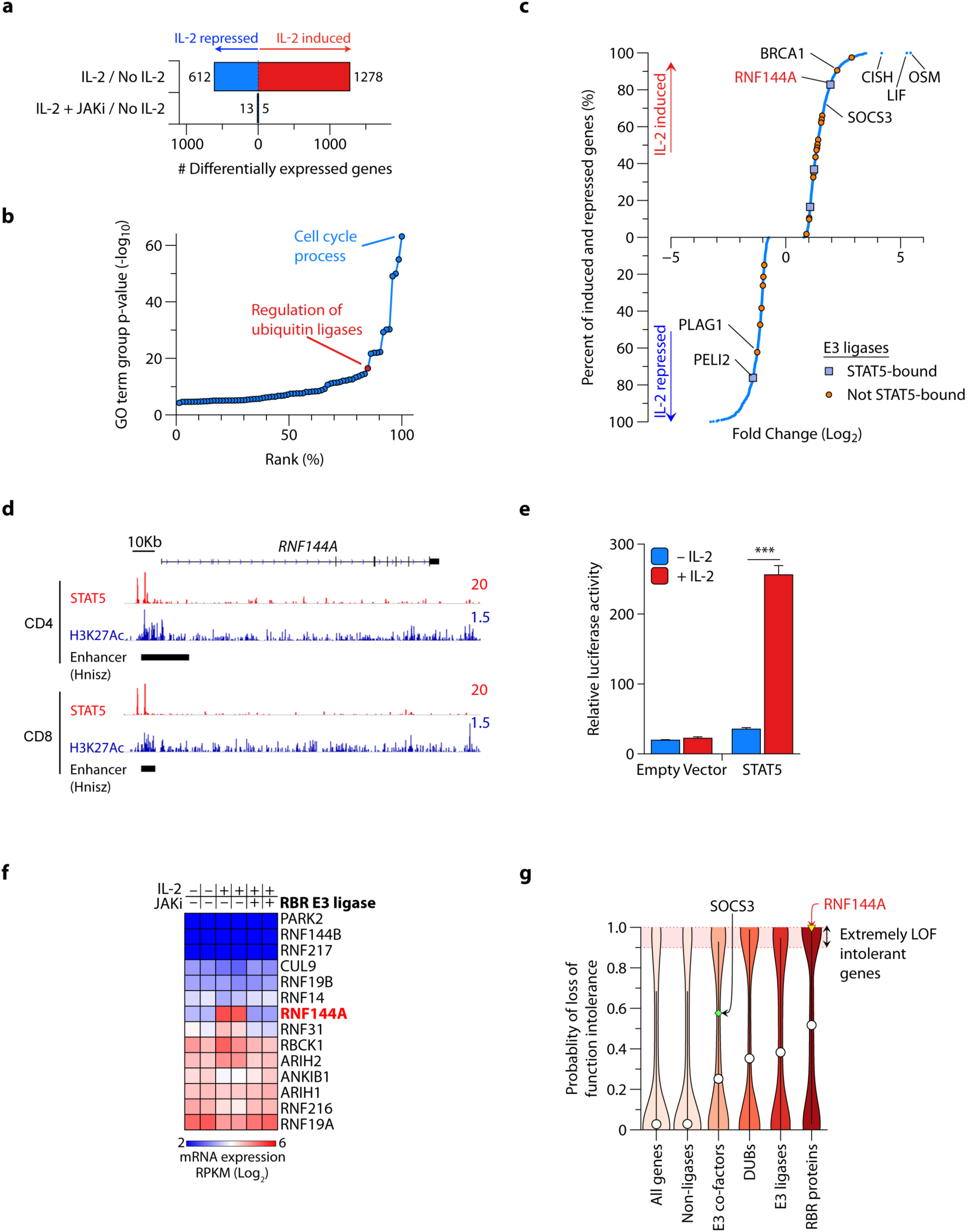
IL-2-induced STAT5 transcriptionally regulates T cell ubiquitination machinery, including the RBR protein RNF144A. **a-b**, Number of differentially expressed genes in CD8^+^ T cells (CTLs) activated, or not, with IL-2 in the presence or absence of a JAK inhibitor (JAKi) (**a**) and ClueGO annotation by Gene Ontology biological process of the 1890 genes differentially regulated by IL-2, ranked by p-value (**b**). **c**, Plot of the 1278 IL-2-induced and 612 IL-2-repressed genes ranked by fold-change; marked are all E3 ligases bound and not bound by STAT5 in ChIP-seq; **a-c** are from *n*=2 replicates. **d**, STAT5 and H3K27Ac ChIP-seq tracks at the *RNF144A* locus in CD4^+^ and CD8^+^ T cells; also shown are enhancer regions annotated in an independent dataset from^62^; shown are representative tracks from *n*=2 replicates. **e**, IL-2 induced STAT5-dependent luciferase reporter activity from a gamma-activated sequence (GAS)-luciferase reporter construct incorporating a GAS motif underlying one of the STAT5 ChIP-seq peaks in (**d**); shown are pooled data from *n*=3 experiments; bars show mean + sem. **f**, Heatmap of gene expression of RBR family members from (**a**). **g**, Violin plots showing distribution of probability of loss of function intolerance (pLI) scores from ExAC for stated families of genes; highlighted are median pLI scores (circles) and pLI scores of two representative genes, *SOCS3* (diamond) and *RNF144A* (triangle). *** p<0.001.

To comprehensively explore this further, we compiled a list of all human E3 ligases (*n*=377), E3 co-factors (*n*=290) and deubiquitinating enzymes (DUBs; *n*=100), as detailed in **methods**, and analyzed the percentage of each group regulated by IL-2 in our dataset and the percentage directly bound by IL-2-induced STAT5 in ChIP-seq. IL-2 regulated approximately 7-10% of each gene group (**Fig. S1b and Table S3**). 15% of E3s (e.g. *RNF157* and *RNF144A*) and 45% of E3 co-factor genes (e.g. *SOCS3* and *CISH*), but none of the DUBs regulated by IL-2 were directly bound by STAT5 (**Figs. S1c, e-f**). Other than ubiquitination, the most common function of IL-2-regulated E3s and E3 co-factors was DNA repair (**Fig. S1d**).

*RNF144A* was the most highly IL-2-induced STAT5-bound E3 ligase gene, falling within the top 20% of all genes induced by IL-2 (**Fig. 1c, Figs. S1e-g**). We confirmed that this gene was IL-2-induced by qPCR (**Fig. S1h**) and that STAT5 bound directly to the gene locus by ChIP-seq (**Fig. 1d**) in both CD4^+^ and CD8^+^ human T cells, with the STAT5 binding sites co-localizing with enhancer marks identified in both cell types in two independent datasets (**Fig. 1d**). Furthermore, IL-2 treatment induced STAT5-dependent luciferase activity in a GAS-luciferase reporter construct incorporating a GAS motif underlying one of the STAT5 ChIP-seq peaks at the *RNF144A* locus, confirming that STAT5 binding induces transcriptional activity (**Fig. 1e**). *Rnf144a* regulation was similar in both mouse and human T cells, although the mRNA peak of *Rnf144a* was earlier in mouse cells (**Figs. S1i**).

RNF144A was the only member of the RBR E3 ligase family significantly induced by the IL-2-JAK/STAT5 pathway (**Fig. 1f**). To contextualize the importance of RNF144A in humans, we sourced estimated probabilities of human gene intolerance to loss of function mutation (pLI scores, where scores of >0.9 indicate extreme intolerance to loss of function mutations) from the ExAC database ^31^. RBR proteins as a family were more intolerant to loss of function than other categories of genes and *RNF144A* was among the most intolerant genes in the entire genome to loss of function, indicating that it is likely to be a critically important gene (**Fig. 1g**). We were unable to find a single *RNF144A* loss of function allele was present in the entire ExAc dataset. Taken together, these results indicate that *RNF144A* is an important *bona fide* IL-2-induced JAK-dependent gene whose transcription is directly regulated by STAT5 in T cells.

### Rnf144a knockout mice are growth-deficient and have abnormal IL-2R signal output

RNF144A is a highly conserved protein containing four domains: RING1, In Between Ring (IBR), RING2 and tail anchor (TA) (**Figs. S2a-d**). We generated a gene knock-out mouse using CRISPR/Cas9 by targeting the first coding exon, resulting in loss of the RING1 domain and a frameshift with a premature translation stop codon in all isoforms of the transcript (**Fig. S3a-c**). *Rnf144a*^*–/–*^ mice were viable, but small for age and growth-deficient (**Fig. 2a and Fig. S3d**). Knockout animals had normal fat and lean body composition making them proportionally small compared to their wild-type littermates (**Fig. S3e**). The cellular composition of the hematological and immunological compartments, including thymus, of unperturbed *Rnf144a*^*–/–*^ mice were unaltered (**Fig. S3f-h**) and these mice did not demonstrate overt evidence of spontaneous autoimmunity.

**Figure 2.**
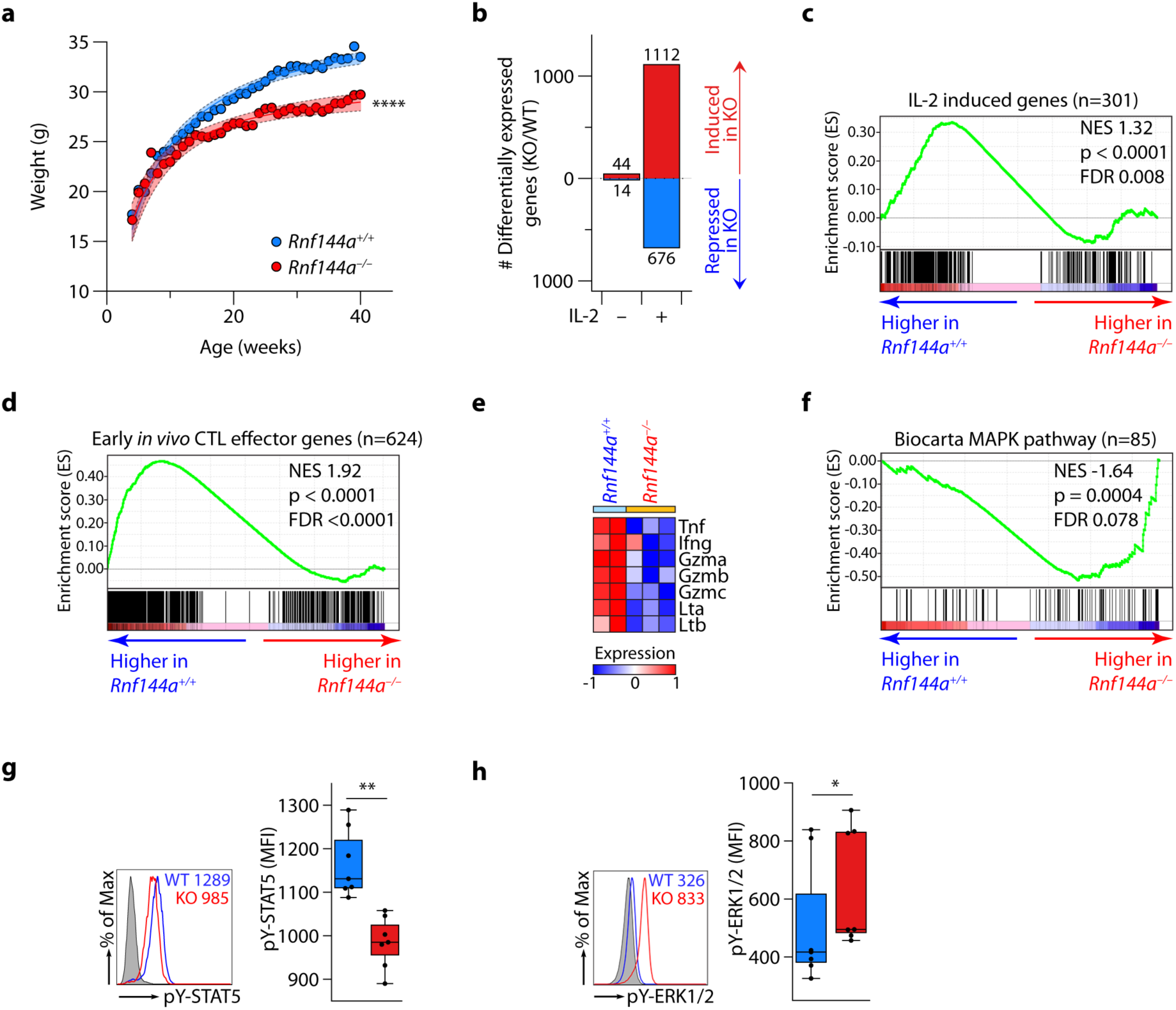
*Rnf144a*^*–/–*^ mice are growth-deficient and have abnormal IL-2R signal output. **a**, Weight gain of *n*=5 littermate *Rnf144a*^*+/+*^ and *Rnf144a*^*–/–*^ mice from weaning to 40 weeks of age. Shown are mean weights (circles) and 95% confidence intervals for the means (dashed lines). **b**, Number of differentially expressed genes in *Rnf144a*^*+/+*^ and *Rnf144a*^*–/–*^ pre-activated CD8^+^ T cells (CTLs) treated, or not, with IL-2; Differentially expressed genes are inferred from *n*=2 *Rnf144a*^*+/+*^ and *n*=3 *Rnf144a*^*–/–*^ mice per group. **c-f**, Geneset enrichment analyses (GSEA) of IL-2-induced (defined as at least 3-fold induction and p<0.05 in wild-type cells on IL-2 treatment) (**c**) and early *in vivo* CTL effector genes (from^33^) (**d**) in IL-2 treated *Rnf144a*^*+/+*^ and *Rnf144a*^*–/–*^ CTLs and heatmap showing expression of key CTL effector genes form the leading edges (**e**). (**f**) GSEA plot of the Biocarta mitogen activated pathway, comparing transcriptomes of IL-2 treated *Rnf144a*^*+/+*^ and *Rnf144a*^*–/–*^ pre-activated CTLs. **g-h**, pY-STAT5 (**g**) and pY-ERK1/2 (**h**) staining in pre-activated primary *Rnf144a*^*+/+*^ and *Rnf144a*^*–/–*^ CD8^+^ T cells in response to IL-2. Shown are representative flow cytometry plots (left) and cumulative data (right) from *n*=3 experiments, respectively. NES = normalized enrichment score; p<0.05; **p<0.01; ****p<0.0001.

RNA-sequencing revealed that, in the absence of IL-2 treatment, pre-activated *Rnf144a*^*–/–*^ CTLs were transcriptionally indistinguishable from those of *Rnf144a*^*+/+*^ mice (**Fig. 2b**). Upon IL-2 stimulation, 1788 transcripts were differentially expressed between the two cell types, indicating dysregulation of IL-2R signal output in knockout cells (**Fig. 2b**). We used Geneset enrichment analysis (GSEA) ^32^ to compare IL-2-treated wild-type and knock-out transcriptomes. We found that wild type (WT) cells were enriched in IL-2-induced transcripts (defined as at least 3-fold induced at p<0.05 in wild-type cells on IL-2 treatment, **Table S5**) (**Fig. 2c**). Similarly, cells lacking *Rnf144a* were enriched in transcripts normally repressed by IL-2 (defined as at least 3-fold induced at p<0.05 in wild-type cells on IL-2 treatment, **Table S5**) (**Fig. S4a**). Importantly, transcripts of IL-2R components were not differently expressed between wild-type and knockout cells (**Figs. S4b-c**). To understand the impact of the altered IL-2-induced transcriptome on CTL function, we sourced early *in vivo* CTL effector genes from the Immunological Genome Project Consortium ^33^ and observed that these genes were enriched in the transcriptomes of WT, compared to *Rnf144a*^−/−^, CTLs (**Fig. 2d**). The core enriched genes in WT cells (**Fig. S4d**) contained those critically important for CTL effector function, including lymphotoxins, granzymes and tumor necrosis factor (TNF) (**Fig. 2e**), suggesting impaired functional fitness of knockout cells.

Unbiased comparisons of the transcriptomes of IL-2-treated WT and *Rnf144a*^*–/–*^ CTLs by GSEA indicated that the genes most differently expressed between the two genotypes were enriched in mitogen activated protein (MAP) kinase signaling, a key pathway downstream of the IL-2R, in knockout cells (**Fig. 2f** and **Figs. S4e-g**), supporting the notion of perturbed transcriptional output downstream of the IL-2R. We hypothesized that the absence of Rnf144a may have perturbed the hierarchy between the STAT5 and ERK-MAPK signaling modules downstream of the IL-2R. We therefore measured activation of STAT5 and ERK following IL-2 treatment in pre-activated wild-type and knockout CTLs. *Rnf144a*^*–/–*^ CTLs had significantly reduced STAT5 and increased ERK phosphorylation in response to IL-2 compared to WT cells (**Figs. 2g-h**). We found no alterations in the expression levels of proximal components of the IL-2R signaling complex, namely IL-2Rβ, IL-2R*γ*, IL-2R*α*, Jak1, Jak3 and STAT5 (**Figs. S4h-m**).

### Rnf144a knockout mice are susceptible to influenza infection

Given that transcriptomes of IL-2-treated *Rnf144a*^*–/–*^ CTLs had suggested impaired cell-intrinsic function (**Figs. 2d-e**), we tested CD8^+^ CTL function *in vivo* using an influenza infection model in which mice were inoculated with sub-lethal doses of murine influenza and culled 8 days later. Parenchymal inflammation following influenza infection was significantly higher in the context of Rnf144a deficiency (**Fig. 3a**). CTLs isolated from draining lymph nodes of influenza infected *Rnf144a*^*–/–*^ mice had a lower frequency of TCR specificity for influenza peptide (**Fig. 3b**) and demonstrated both impaired cytotoxicity (reduced expression of CD107a/b) and effector cytokine production, including TNF and IFN*γ*, in response to re-challenge with influenza peptide (**Fig. 3c**). Furthermore, antigen-specific tissue resident memory cells were reduced in number in the lungs of influenza-infected mice harvested 30 days after infection (**Fig. S5a**). These data confirmed that *Rnf144a* deficiency results in impaired anti-viral immunity.

**Figure 3.**
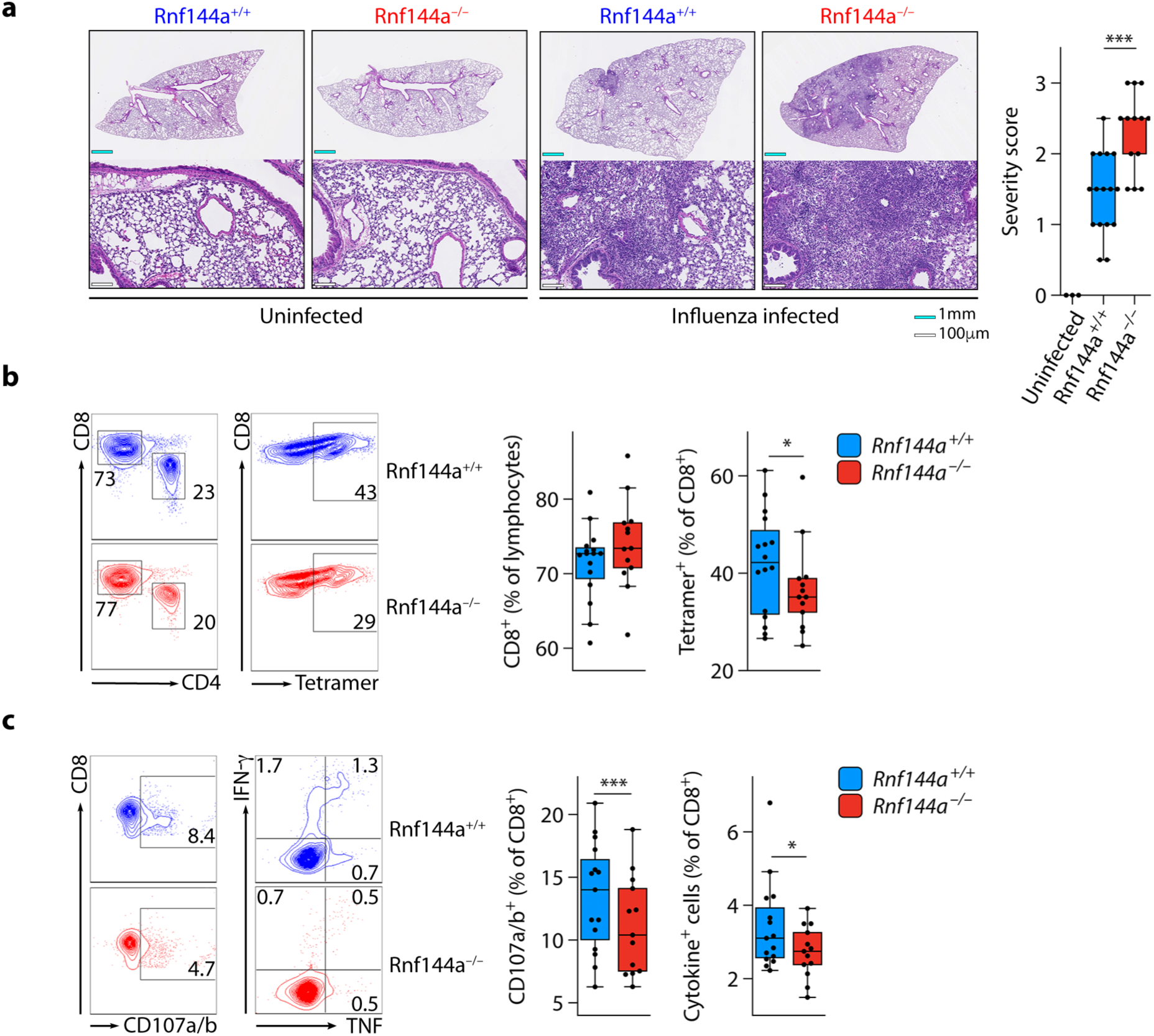
*Rnf144a*^*–/–*^ mice are susceptible to influenza infection. **a**, H&E-stained day eight lung histology from tissues of *Rnf144a*^*+/+*^ and *Rnf144a*^*–/–*^ mice infected, or not, with sub-lethal doses of influenza; shown are representative histology slides (left) and cumulative severity scores (right). **b**, Proportions of CD8^+^ T cells (CTLs) and percentage of antigen-specific CTLs in tissues of influenza-infected *Rnf144a*^*+/+*^ and *Rnf144a*^*–/–*^ mice harvested on day 8. **c**, Degranulation (CD107a /b expression) and cytokine production (IFN*γ* and TNF) of CTLs from influenza-infected *Rnf144a*^*+/+*^ and *Rnf144a*^*–/–*^ mice re-challenged *in vitro* with influenza peptide. **a-c** show representative examples and cumulative data from *n*=2 independent experiments. *p<0.05 **p<0.01 ***p<0.001.

### RNF144A enhances JAK-STAT5 activation by promoting STAT5 interaction with IL-2Rβ

RNF144A is one of only three members of the RBR proteins with a C-terminal membrane tail-anchor (TA) domain ^34^ (**Fig. S2a**). We confirmed the requirement of the TA for plasma membrane localization of RNF144A using membrane flotation sucrose gradients assays and immunofluorescence studies and, importantly, established that the RING domains of RNF144A are cytoplasmically oriented, allowing for interactions with intracellular proteins (**Figs. S6a-d**). We also confirmed the ability of RNF144A to behave as a RING-HECT E3-ubiquitin ligase, as it can function with the E2 protein UbcH7 and other E2s like UbcH5a (**Figs. S7a-c**).

As RNF144A is induced by IL-2 and localized to the plasma membrane, we suspected that potential substrate(s) of RNF144A might be membrane proximal components of the IL-2 signaling machinery. To confirm this, we generated a specific knockdown of RNF144A in the IL-2-responsive cell line, YT, using lentiviral delivery of RNF144A-specific shRNA (“KD” cells) or a control non-specific shRNA (“Control” cells) (**Fig. 4a**). Absence of RNF144A impeded the known IL-2-dependent cellular functions of these cells, notably IFN*γ* production ^35^ and cytotoxicity ^36^ (**Figs. S8a-b**). Furthermore, Western blot analysis of global IL-2-induced tyrosine phosphorylation (pY99) showed clear differences in the kinetics and magnitude of phosphorylation for a number of proteins in KD versus control cells (**Figs. S8c**), suggesting altered IL-2 responsiveness of cells deficient in RNF144A, similar to the observations from *Rnf144a*^*–/–*^ mice. KD cells had decreased IL-2-induced STAT5 activation (**Fig. 4b**, quantified in **Figs. S8d**). Furthermore, RNF144A overexpression in an IL-2R reconstitution system ^37^ enhanced IL-2-induced STAT5 phosphorylation (**Fig. 4c**), as well as STAT5-dependent transcriptional activation of a luciferase reporter gene (**Fig. 4d**). Thus, RNF144A is a positive regulator of STAT5 signaling.

**Figure 4.**
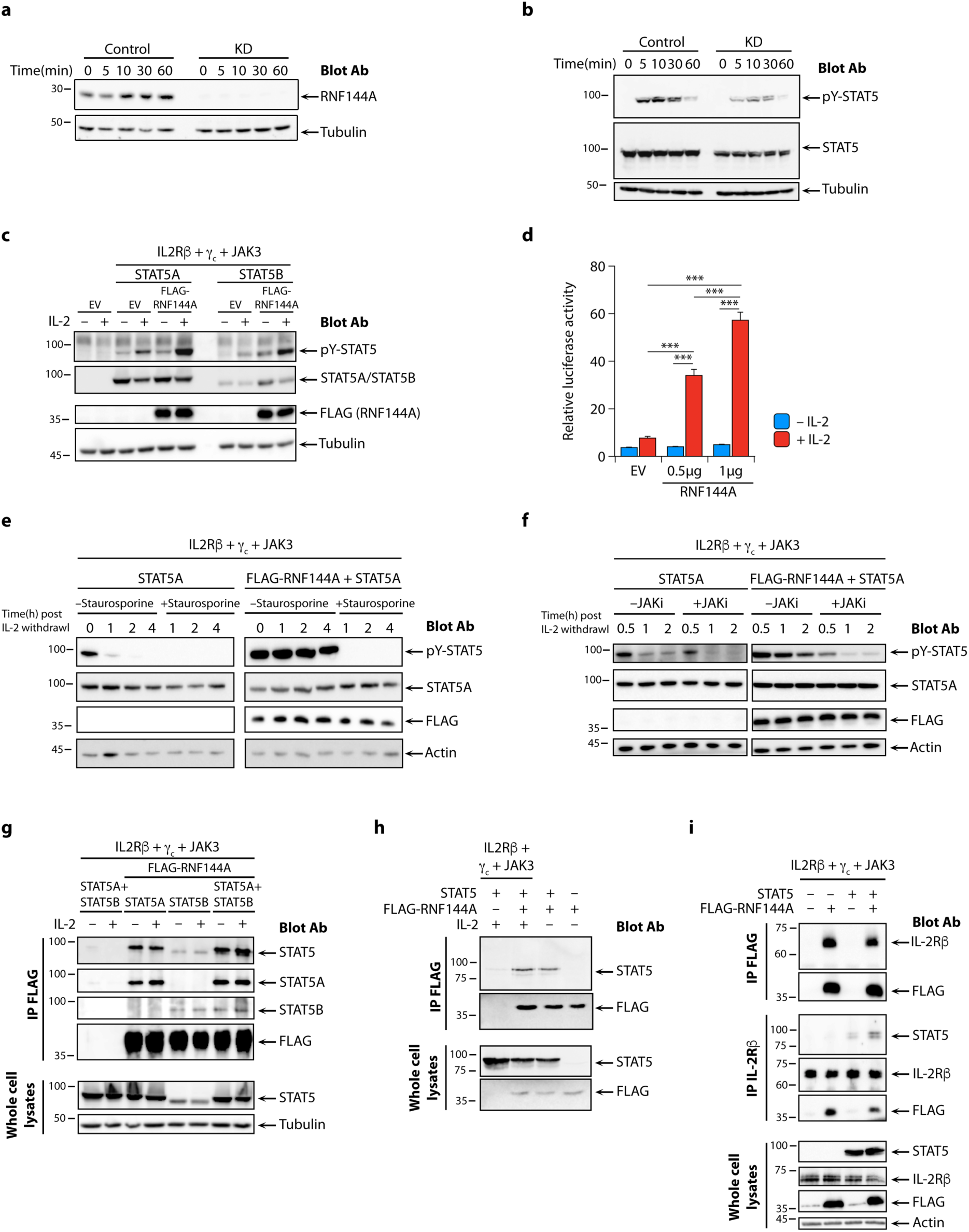
RNF144A regulates IL-2R-STAT5 signal transduction. (**a**) Western blot of RNF144A expression following stimulation with IL-2 in whole cell extracts of YT cells stably transduced with non-specific shRNA (Control) or shRNA targeting RNF144A for knockdown (KD); (**b**) Western blots showing pY-STAT5 phosphorylation in whole cell lysates of IL-2 stimulated Control and KD cells (quantified in **Supplementary Fig. 8d**); (**c**) Western blots showing IL-2-induced pY-STAT5 phosphorylation in HEK293T cells transfected with IL-2R*β, γ*c, JAK3, STAT5A or STAT5B with or without FLAG-RNF144A; (**d**) IL-2 induced STAT5-dependent luciferase reporter activity in HEK293T cells transfected with IL-2R*β, γ*c, JAK3, STAT5A and either empty vector (EV) or RNF144A; (**e-f**) Western blots showing pY-STAT5 phosphorylation after IL-2 withdrawal in the presence or absence of staurosporine (**e**) or a JAK inhibitor (JAKi) (**f**) in HEK293T cells transfected with IL-2R*β, γ*c, JAK3, STAT5A with or without FLAG-RNF144A; (**g**) Western blots for STAT5, STAT5A and STAT5B from immunoprecipitates of FLAG or whole cell lysates of HEK293T cells transfected with IL-2R*β, γ*c, JAK3, STAT5A and/or STAT5B with or without FLAG-RNF144A. Note that STAT5A-specific antibodies for Western blotting are more robust than STAT5B-specific antibodies; (**h**) Western blots for STAT5 from immunoprecipitates of FLAG or whole cell lysates of HEK293T cells transfected, as indicated, with IL-2R*β, γ*c, JAK3, and/or STAT5 with or without FLAG-RNF144A; (**i**) Western blots for STAT5, IL-2Rβ, and FLAG of anti-FLAG or anti-IL-2Rβ immunoprecipitates from unstimulated whole cell lysates of HEK293T cells transfected with IL-2R*β, γ*c, JAK3 with or without STAT5 and/or FLAG-RNF144A. Control whole cell lysate blots to detect STAT5, IL-2Rβ, FLAG and actin are also shown (quantified in **Supplementary Fig. 8f**). Shown are representative examples from multiple experiments (minimum *n*=3). Bars are mean + sem; ***p<0.001.

The increased STAT5 activation in the presence of RNF144A could either be due to increased phosphorylation by JAKs or delayed dephosphorylation of phospho-STAT5. To distinguish between these possibilities, we first measured the kinetics of STAT5 de-phosphorylation upon IL-2 withdrawal in the presence or absence of RNF144A. Without RNF144A, STAT5 was dephosphorylated within 1 hour, while in the presence of RNF144A phosphorylation of STAT5 was sustained for even 4hrs after IL-2 withdrawal (**Fig. 4e**). IL-2 withdrawal in the presence of the tyrosine kinase inhibitor staurosporine, or a JAK-specific inhibitor, resulted in immediate abrogation of STAT5 phosphorylation, even in RNF144A-overexpressing cells (**Figs. 4e-f**), indicating that RNF144A functions primarily by promoting the continued phosphorylation of STAT5 by JAKs. RNF144A and STAT5 co-localized at the plasma membrane (**Fig. S8e**) and a physical interaction between the two was confirmed by co-immunoprecipitation of both STAT5A and STAT5B with RNF144A in the presence or absence of IL-2R (**Figs. 4g-h**). Furthermore, RNF144A and IL-2Rβ were co-immunoprecipitated with either anti-IL-2Rβ or anti-RNF144A (FLAG) antibodies and, strikingly, co-precipitation of STAT5 with IL2-Rβ was significantly increased in the presence of RNF144A (**Fig. 4i;** quantified in **Fig. S8f**). These data indicate that RNF144A exists in complex with STAT5 and IL-2Rβ, enhancing STAT5-IL-2Rβ interaction, prolonging JAK-mediated STAT5 activation and STAT5-transcriptional output from the IL-2R.

### RNF144A restricts IL-2R-ERK MAPK signaling by ubiquitination of RAF1

We next investigated how the absence of RNF144A enhances ERK signaling from the IL-2R (**Fig. 2h**). Deficiency of RNF144A in KD cells significantly enhanced and sustained ERK1/2 activation compared to control cells (**Fig. 5a** and **Fig. S9a**). As total ERK protein expression was similar in these cells, we studied the other upstream activating kinases in this pathway. Phosphorylation of RAF1 was significantly enhanced in IL-2 treated KD cells and the expression of total RAF1 protein was basally increased in cells lacking RNF144A (**Fig. 5b**; quantified in **Fig. S9b**). Serial dilutions of cell lysates from unstimulated cells confirmed that homeostatic levels of RAF1 were increased by approximately 2-3-fold in cells lacking RNF144A (**Fig. 5c**; quantified in **Fig. S9c**). RAF1 can heterodimerize with BRAF to form a potent ERK activating complex^60^. We speculated that the increased RAF1 expression in KD cells may impact on RAF1-BRAF heterodimer formation. Substantially more BRAF co-immunoprecipitated with RAF1 following IL-2 treatment in the absence of RNF144A (**Fig. 5d**), indicating that RNF144A restrains ERK activation by inhibiting heterodimerization of RAF1 with BRAF.

**Figure 5.**
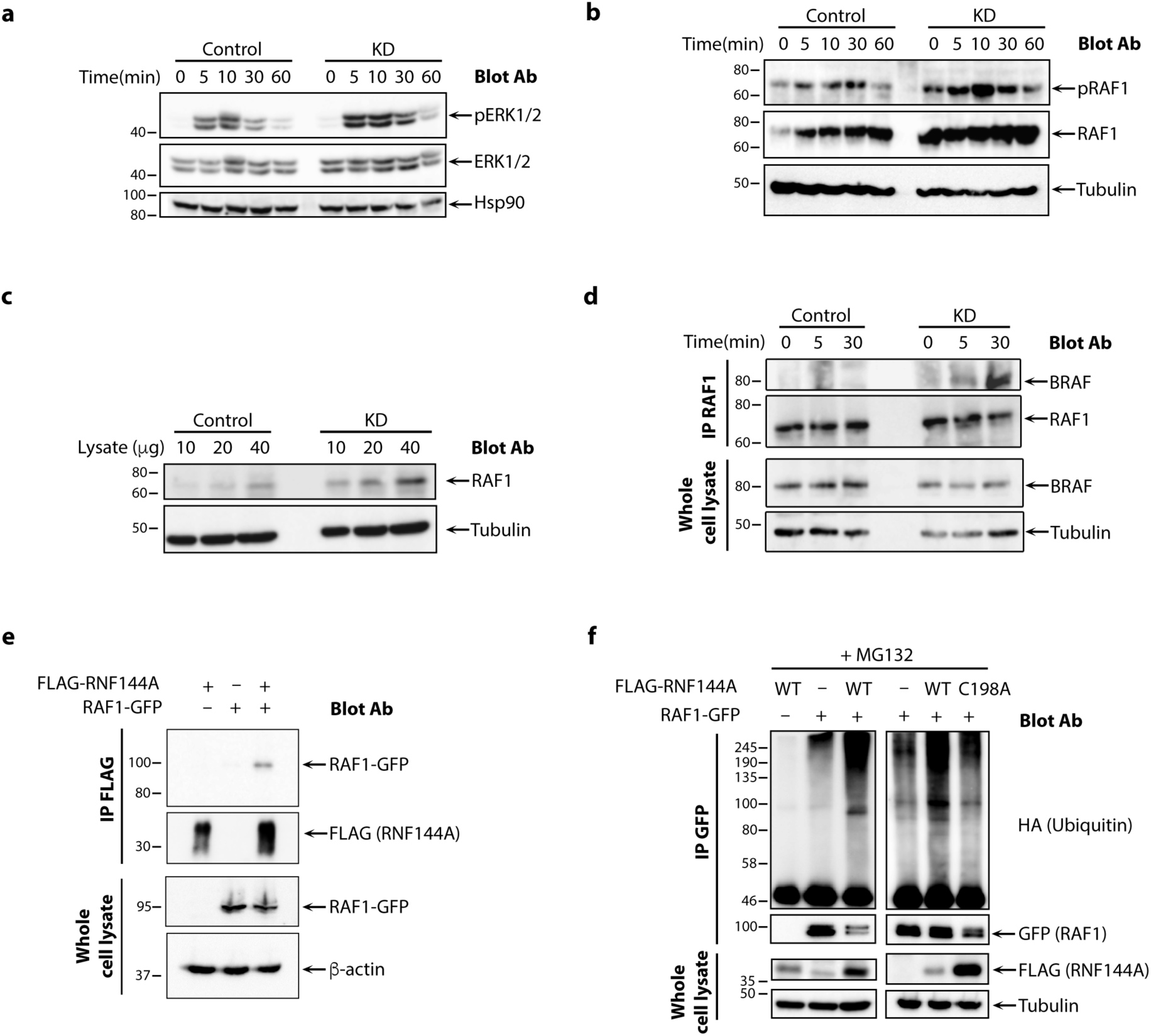
RNF144A directly regulates the MAPK-ERK pathway by polyubiquitination of RAF1. **a-b**, Western blots showing total and phosphorylated ERK (**a**) and total and phosphorylated RAF1 (**b**) in Control and KD cells after treatment with IL-2 (time zero quantified in **Supplementary Fig. 9b**). **c**, Western blots showing homeostatic levels of total RAF1 in serial dilutions of cell lysates from Control and KD cells (quantified in **Supplementary Fig. 9c**). **d**. Western blots for RAF1 and BRAF from immunoprecipitates of RAF1 or whole cell lysates from Control and KD cells treated with IL-2. **e**, Western blots for RAF1 and FLAG from immunoprecipitates of FLAG and whole cell lysates in HEK293T cells transfected with RAF1-GFP and FLAG-RNF144A. **f**, HA blots from MG132-treated whole cell lysates and GFP immunoprecipitates of HEK293T cells transfected with HA-ubiquitin and combinations, as indicated, of wild type (WT) or C198A mutant FLAG-RNF144A and RAF1-GFP. Western blots throughout are representative examples of at least *n*=3 experiments each.

Increased basal RAF1 in the absence of RNF144A suggested that RAF1 may be a substrate for RNF144A-mediated ubiquitination. RNF144A co-localized with RAF1 in immunofluorescence studies of transfected HEK293T cells (**Fig. S9d**). Furthermore, RAF1 co-immunoprecipitated with RNF144A, indicating that the two proteins exist in a complex *in vivo* (**Fig. 5e**). To determine whether RNF144A regulates RAF1 ubiquitination, we assayed the ubiquitination of RAF1-GFP by RNF144A *in vivo* in transfected HEK293T cells. WT and RING/HECT-inactive mutant of RNF144A (C198A) were overexpressed in HEK293T cells along with HA-ubiquitin (HA-ub) and ubiquitination of RAF1 was investigated. Co-expression of WT RNF144A, but not RNF144A^C198A^, resulted in increased polyubiquitination of RAF1, detected in the presence of the proteasome inhibitor, MG132 (**Fig. 5f**). Conversely, Western blot analysis of RAF1 immunoprecipitates from control and KD cells revealed a substantial decrease in IL-2-induced ubiquitinated RAF1 in KD cells, despite more RAF1 immunoprecipitation, compared to controls (**Fig. S9e**). Taken together, these data confirm that RAF1 is a substrate for RNF144A-mediated ubiquitination and proteasomal degradation.

### Reduced RNF144A expression is associated with greater severity of human influenza infection

The impaired anti-viral immunity to influenza and greater tissue inflammation in the context of murine *Rnf144a* deficiency suggested that RNF144A may also be associated with clinical influenza severity in humans. To explore this possibility, we examined whole blood transcriptomes from a large dataset of healthy subjects and patients with confirmed moderate or severe influenza (GSE101702 ^38^) for expression of *RNF144A*. mRNA encoding RNF144A was significantly lower in patients with severe influenza compared to healthy controls or those with moderate disease (**Fig. 6a**) and ERK-regulated genes were significantly enriched in transcriptomes of patients with severe influenza compared to both other subject cohorts (**Figs. 6b-c**). The leading edge (core enriched) of ERK-regulated genes clustered with disease severity (**Fig. 6d**) and strongly correlated positively with each other across all donors, suggesting that they are co-regulated (**Fig. 6e**). Conversely, these genes were significantly negatively correlated with the expression of *RNF144A* (**Figs. 6e-f**). We tested the performance of *RNF144A* as a biomarker to distinguish severe influenza from moderate influenza, together with *FOS* and *JUN*, which are AP-1 subunits and downstream targets of ERK. The receiver operating characteristic (ROC) curve for *RNF144A* was highly significant (AUC = 0.71; *p* = 0.0002) and performed well in comparison with *JUN* (AUC = 0.76; *p* <0.0001) and *FOS*, which was not a predictor (AUC = 0.56; *p* = 0.28) (**Fig. 6g**), when predicting severe versus moderate influenza. We next compared the performance of *RNF144A* against those of published genes associated with influenza severity, *IFITM3* ^39^, *TLR3* ^40^, *MX1* (in mice) ^41^, *IRF7* ^42^, *GATA2* ^43^ and *CD177* (a neutrophil-specific marker) ^38^, as well as *CD8A*. Only *CD177, IFITM3, TLR3* and *RNF144A* expressions were able to predict influenza severity. Of these, *CD177* and *RNF144A* were the best performing predictors (**Fig. 6h**). These data indicate that *RNF144A* expression in human whole blood transcriptomes is negatively associated with influenza severity and with expression of ERK target genes and performs better than most other available biomarkers to distinguish severe from moderate influenza.

**Figure 6.**
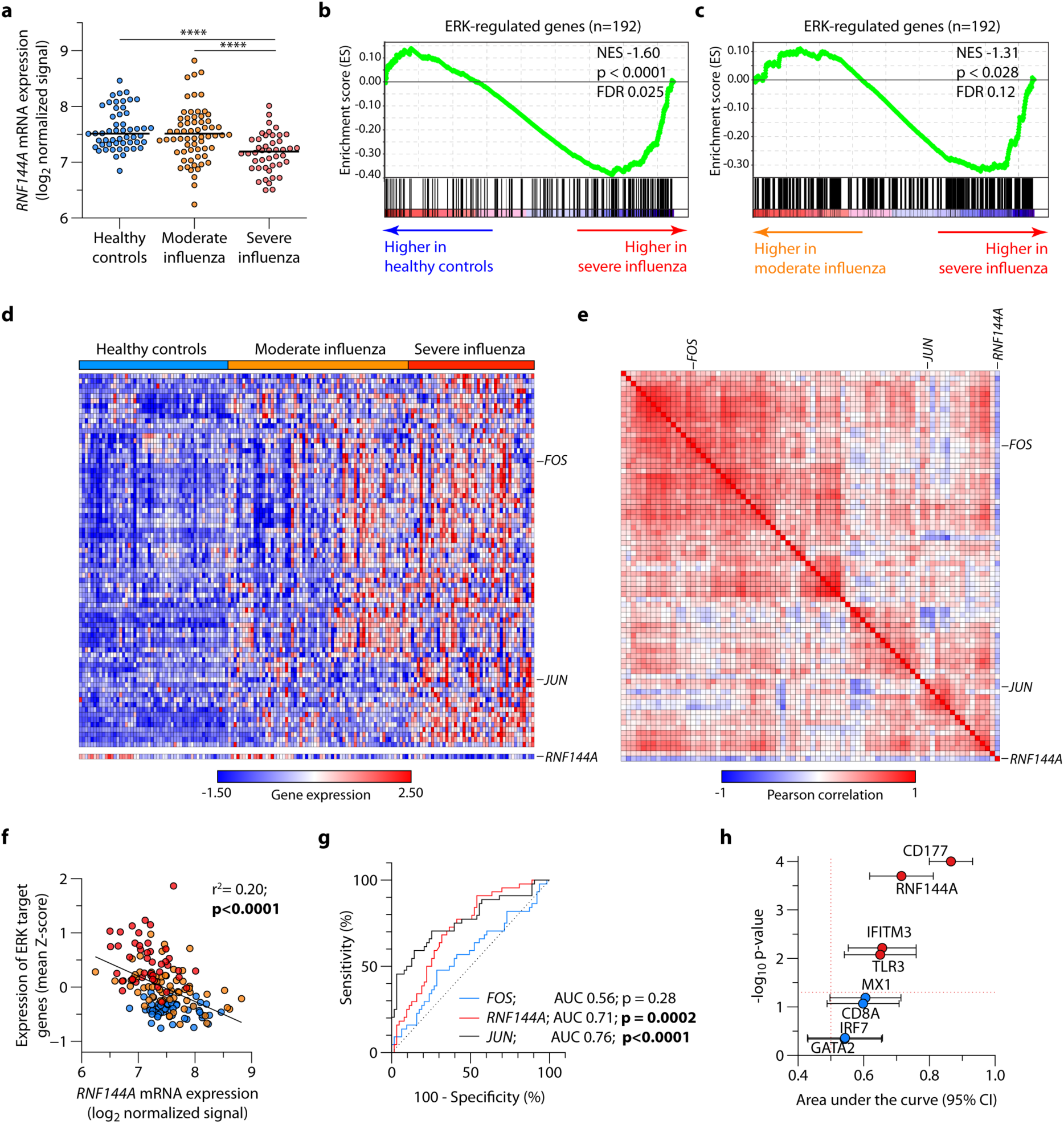
RNF144A expression is a biomarker of influenza severity in patients. **a**, Expression of RNF144A mRNA in healthy control PBMC and PBMCs from patients with moderate and severe influenza. **b-c**, Geneset enrichment analysis (GSEA) of ERK-regulated genes comparing patients with severe influenza to healthy controls (**b**) and to patients with moderate influenza (**c**). **d**, Expression of the core (leading edge) enriched genes from **b** in healthy controls and patients with influenza. Shown, in addition, is expression of *RNF144A* mRNA. **e**, Correlation matrix of genes in (**d**) across all samples. **f**, Linear regression line between *RNF144A* mRNA and mean expression (Z-score) of leading edge ERK-regulated genes. Each point indicates one subject, colored by phenotype (blue = healthy control, orange = moderate influenza, red = severe influenza); indicated is the Pearson correlation co-efficient and its p-value. **g**, ROC curves showing performance of *FOS, JUN* and *RNF144A* mRNA expression to discriminate moderate from severe influenza. AUC analyses and their p-values indicated. **h**, P-values and AUC + 95% confidence interval (CI) of ROCs for indicated transcripts to distinguish moderate from severe influenza. Source data for (**a-h**): GSE101702^38^; *n*=52 healthy donors, *n*=63 with moderate influenza and *n*=44 with severe influenza throughout. NES = normalized enrichment score ****p<0.0001.

## Discussion

STAT5 proteins are key signaling mediators facilitating the unique transcriptional response downstream of IL2/IL-2R, but the molecular mechanisms ensuring dominance of the JAK/STAT5 pathway over the concurrently activated RAS-MAPK and PI3K-AKT pathways are poorly understood. Here we set out to identify novel mechanisms and regulators of IL-2 signaling that promote STAT5 activation. We identified an unexpected role for the E3-ubiquitin ligase RNF144A in the regulation of IL-2 signaling. Using a combination of transcriptomic, biochemical, and genetic approaches we demonstrated that *RNF144A* is an IL-2 induced STAT5-regulated gene that acts in a feedback loop to enhance STAT5 activation while simultaneously restraining MAPK-ERK signaling by targeting RAF1 for proteasomal degradation. The physiological importance of Rnf144a is underscored by the conspicuous immunodeficiency of *Rnf144a*^*–/–*^ mice manifest upon infection.

A striking result from RNA-seq was the incidence of genes associated with ubiquitination processes, occurring within the top 15% of all functional gene categories induced by IL-2, suggesting an important role for ubiquitination in shaping the downstream biological functions of this cytokine. With the exception of UHRF1 ^44^, the regulation of the E3s identified in our study have not been previously linked to IL-2, including well known ligases such as BRCA1 and PELI2. Only a subset of these, including RNF144A, but excluding UHRF1, were directly regulated by STAT5 in ChIP-seq studies. To our knowledge, this is the first comprehensive delineation of transcriptional regulation of the ubiquitination machinery downstream of IL-2 in CTLs. Interestingly, many of the E3-ligases identified as IL-2-regulated in this analysis, have functions associated with DNA repair. Given that IL-2 promotes clonal expansion of activated T cells, involving rapid DNA synthesis and replication ^45^, the finding that IL-2 induces an ubiquitome dedicated to DNA repair is consistent with this. Indeed, somatic mutations in *RNF144A* have been identified in several cancer studies ^27^, despite the absence of germline loss of function mutations in whole exome databases.

Furthermore, RNF144A expression is dysregulated in autoimmune diseases, such as Wegener’s granulomatosis, malignancies, such as esophageal cancer, renal carcinoma and T cell lymphomas, and genome wide association studies suggest an association between single nucleotide variants of *RNF144A* and neurological and inflammatory diseases ^46-49^, indicating that RNF144A, like other members of the RBR family, plays important physiological roles in the homeostatic regulation of both immune and non-immune cell functions. In comparison to the other RBR family proteins, including the closest related RBR family member, RNF144B, a regulator of cell proliferation and homeostasis ^50,51^, RNF144A was the only IL-2-induced JAK-dependent and STAT5-bound gene, indicating specificity and non-redundancy between these two proteins.

Surprisingly, despite the prediction from ExAC that humans are highly intolerant to loss of RNF144A, *Rnf144a*^*–/–*^ mice were viable. *Rnf144a*^−/−^ mice displayed no gross phenotypes under sterile homeostatic conditions apart from runted appearance, reminiscent of male *STAT5b*^*–/–*^ mice, which display growth retardation due to defects in growth hormone-induced STAT5 signaling ^52^, suggesting a STAT5 activation deficiency in *Rnf144a*^*–/–*^ mice, that was not limited to T cells. Members of the RBR family each have multiple substrates (Parkin, for example, has been estimated to have between 90-1700 protein substrates ^34^), thus it is highly likely that RNF144A also has multiple biological functions *in vivo*. In this study, we have limited our investigations to focus primarily on IL-2-JAK/STAT5 dependent responses in CTLs, where IL-2-stimulated STAT5 activation was significantly decreased and ERK activation increased in *Rnf144a*^*–/–*^ cells, mirroring RNF144A-knockdown cells. It should be noted that STAT5 activation in *Rnf144a*^*–/–*^ mouse cells was decreased but not absent. Thus, classical Stat5 deficiency phenotypes, such as reduction in nTregs and spontaneous autoimmunity, was consistently not evident under sterile conditions as many of these phenotypes require complete deletion of at least three out of four alleles of *STAT5* and a low threshold of IL-2R/STAT5 signaling is sufficient to promote nTreg development and function ^12,53^. Furthermore, the impaired STAT5 signaling may be sufficient to maintain homeostasis until the immune system is stressed, i.e. during an infection.

Impaired STAT5 signal output from the IL-2R in the absence of Rnf144a was suggested by functional data revealing immunodeficiencies in *Rnf144a*^−/−^ mice associated with reduced IL-2-induced STAT5 activity. As effector functions in CTLs are IL-2-dependent ^34^, the manifest phenotype in response to influenza infection was extensive tissue inflammation, impaired cytotoxicity and effector cytokine production. Similar observations in human cohorts suggest the possibility that RNF144A expression could act as a biomarker to distinguish severe from moderate influenza and/or that modulating RNF144A function biochemically could have therapeutic benefit for patients hospitalized with severe influenza. Of note, earlier studies on viral replication and nuclear export mechanisms have shown that influenza A infection leads to activation of the Raf/MEK/ERK signaling cascade and that specific inhibitors of this pathway can effectively block virus propagation ^54^. Our observations that loss or reduction of RNF144A results in impaired anti-viral immunity to influenza in both mice and man, suggest that RNF144A is an essential cellular inhibitor of the RAF-ERK-MAPK pathway, required to prevent pathogenic anti-viral responses by CTL.

Silencing of RNF144A in the IL-2-inducible cell line YT provided a useful tool to interrogate pathways and substrates targeted by RNF144A after IL-2 stimulation. Functional studies indicated a role for RNF144A in regulation of IL-2-mediated signal transduction and downstream effector functions, such as cytotoxicity and IFN*γ* production, which are JAK3-STAT5 pathway dependent ^2^. In keeping with this, IL-2-induced STAT5 activation was decreased in RNF144A-deficient cells. There was both a physical and functional relationship between RNF144A and STAT5, with the E3 sustaining both duration and magnitude of IL-2-activated STAT5 and existing in complex with STAT5 and the IL-2Rβ. Surprisingly, we found no *in vitro* evidence for direct ubiquitination of STAT5 by RNF144A (unpublished observations), despite detecting physical interactions between the two proteins, suggesting that RNF144A regulates STAT5 indirectly via modulation of another substrate, which affects STAT5 function or by acting as a platform to enhance physical association between STAT5 and the IL-2R. While our biochemical data provided evidence to support the latter, the detailed mechanism(s) by which RNF144A enhances STAT5 activation requires further clarification in future studies.

Concomitant with a decrease in IL-2-induced genes, there was an unexpected significant activation of ERK-induced genes in the absence of RNF144A, indicative of a direct impact of RNF144A on one or more components of the ERK-MAPK pathway, downstream of IL-2R. We identified physical and functional interactions between RNF144A and RAF1, resulting in the polyubiquitination and targeting of RAF1 for proteasomal degradation. RAF1 is a notable candidate for RNF144A regulation as it is constitutively associated with IL-2R prior to IL-2 stimulation and its expression and kinase activity are induced by IL-2, leading to downstream activation of ERK ^55,56^. RAF1 can be targeted for proteasomal degradation by CHIP- and CTLH-mediated ubiquitination in non-immune cells ^57,58^. The relevance of these pathways have not been confirmed in immune cells and neither CHIP nor CTLH were induced by IL-2 in our gene expression studies. We believe that this is the first demonstration of a cytokine-induced E3 that regulates homeostatic RAF1 levels in immune cells, potentially an important observation as dysregulated RAF1 is oncogenic.

RAF1 can dimerize with itself or with other RAF family members, e.g. BRAF; the heterodimer formed between RAF1 and BRAF has substantially increased kinase activity compared to the respective monomers or homodimers and is cancer associated ^59,60^. RNF144A restricted not only RAF1 activation, but perhaps more importantly, formation of RAF1/BRAF heterodimers both basally and after IL-2R engagement. Increased RAF1/BRAF significantly impacts downstream ERK activation ^60^. Thus, our studies reveal an unexpected critical role for RNF144A in orchestrating “faithful” IL-2R signaling to qualitatively fine-tune the transcriptional response by promoting STAT5 activity, while limiting RAF-ERK activity.

As this study focused primarily on identifying regulators of the JAK-STAT5 pathway, it remains to be established if RNF144A has any impact on the JAK-independent, PI3K pathway downstream of the IL-2R. Of note, DNA-PKcs, an established substrate of RNF144A downstream of growth factor receptors ^27^, can phosphorylate Ser-473 of AKT to activate it during the physiological response to DNA-damage ^61^. However, the role of DNA-PKcs (encoded by *PRKDC*) in IL-2R function is unknown and remains to be determined in future studies.

In conclusion, RNF144A is an IL-2-induced, STAT5-regulated, membrane-associated E3 ubiquitin ligase that fine-tunes IL-2R signaling output by restraining the RAF-ERK-MAPK pathway via regulation of RAF1 levels and allows rapid STAT5 activation to proceed unperturbed. Taken together, these data indicate that RNF144A is a critical component of healthy immunity and that its absence causes immunodeficiency, manifest as more severe viral infection.

## Acknowledgements

This work was supported by the Medical Research Council (G0400197 to S.J.), Wellcome Trust (grant 097261/Z/11/Z to B.A., 102932/Z/13/Z to C.K.), the Crohn’s & Colitis Foundation of America CCFA (A.L.), the Purdue Cancer Center (M.K.), the Wegener’s Trust (S.K. and S.J.) and National Heart, Lung, and Blood Institute (grant 5K22HL125593 to M.K.). Research was also supported by the National Institute for Health Research (NIHR) Biomedical Research Centre at Guy’s and St Thomas’ NHS Foundation Trust and King’s College London. The views expressed are those of the author(s) and not necessarily those of the NHS, the NIHR or the Department of Health. This research was supported in part by the Intramural Research Programs of the National Institute of Arthritis and Musculoskeletal and Skin Diseases, the National Institute of Allergy and Infectious Diseases, the National Institute of Diabetes and Digestive and Kidney Diseases of the National Institutes of Health. We thank Matt Arno (Genomics Centre, King’s College London) with gene expression microarray studies. In addition, the authors thank Adrian Hayday (CRICK-KCL, UK), Sarah Gaffen (Pittsburgh University), Helen Collins (KCL, UK), Han-Yu Shih (NIH) and Fred Davis (NIH) for critically reading the manuscript.

## Author contributions

B.A. and S.J. designed and performed experiments, analyzed data and wrote the manuscript. J.J.O’S. provided scientific input, supervised the project and wrote the manuscript. S.J. originated and conceptualized the study, supervised the project and wrote the manuscript. All other authors performed experiments, analyzed data and/or provided scientific input.

## Supplementary information

**Supplementary Figs 1-9 uploaded as a separate file**

**Supplementary Tables**

**Supplementary Table 1. List of differentially expressed genes in CTLs**.

**Supplementary Table 2. Functionally grouped terms in ClueGO**.

**Supplementary Table 3. List of IL-2 regulated E3 ligases, E3 co-factors and DUBs and STAT5-binding**.

**Supplementary Table 4. List of human E3 ligases, E3 co-factors and DUBs.**

**Supplementary Table 5. Genesets used in GSEA**.

## Materials and Methods

### Ethics approvals

All animal studies were performed according to National Institutes of Health guidelines for the use and care of live mice and were approved by the Institutional Animal Care and Use Committee of National Institute of Arthritis, Musculoskeletal and Skin Diseases (Protocol number A017-02-01) and National Institute of Diabetes and Digestive and Kidney Diseases (Protocol number K051-KDB-18).

### Antibodies and other reagents

The following anti-human protein antibodies were used in these studies: anti-pan-STAT5 (BD Biosciences), anti-STAT5a and -STAT5b specific antibodies (SantaCruz Biotechnologies), anti-pY694-STAT5 (New England Biolabs), anti-RNF144A (Acris), anti-tubulin (Sigma), anti-pTyr (pY99) (SantaCruz Biotechnology), anti-Hsp90 (SantaCruz Biotechnology), anti-actin (SantaCruz Biotechnology), anti-pan-ERK (Cell Signaling), anti-p44/42ERK1/2 (Cell Signaling), anti-RAF1 (SantaCruz Biotechnology), anti-pS338-RAF1 (Cell Signaling), anti-BRAF (SantaCruz Biotechnology), anti-ubiquitin VU-1 (Tebu-bio), anti-HA (Covance), anti-transferrin receptor (Zymed), anti-p23 (Alexis Biochemicals), anti-FLAG M2 monoclonal (Sigma), anti-GST (Amersham) and anti-GM130 (Abcam), anti-GAPDH (Proteintech). In addition, the following secondary reagents were used: goat anti-mouse HRP (Perbio), donkey anti-mouse HRP (Perbio), donkey anti-mouse-HRP (GE Healthcare), donkey anti-rabbit-HRP (GE Healthcare), goat anti-mouse AlexaFluor594 (Molecular Probes), streptavidin-HRP (Perbio), anti-goat HRP (SantaCruz Biotechnology), anti-rabbit HRP (LiCor). MitotrackerRed850 was purchased from Molecular Probes. RAF1-GFP was generous gift from Dr. Walter Kolch (Systems Biology Ireland & Conway Institute University College Dublin, Ireland).

The following anti-mouse antibodies were used for flow cytometry in these studies: anti-mouse CD3 (500A2), anti-mouse CD4 (RM4-5), anti-mouse CD8 (56-6.7), anti-mouse CD16/CD32 (clone 2.4G2), anti-mouse CD44 (IM7), anti-mouse CD49b (DX5), anti-mouse CD69 (H1.2F3), anti-mouse CD122 (TM-Beta 1), anti-mouse CD138 (281-2), anti-mouse CXCR5 (2G8), anti-mouse Fas (15A7), anti-mouse IFN*γ* (XMG1.2), anti-pT202/pY204 ERK1/2 (20A), pY694-Stat5 (47/Stat5(Y694)), anti-mouse B220 (RA3-6B2), streptavidin-APC (all BD), anti-mouse CD25 (BC61.5), anti-mouse GL7 (GL-7), anti-mouse NKp46 (29AI.4), anti-mouse IgD (11-26), anti-mouse PD1 (J43), anti-mouse Foxp3 (FJK-16s) (all eBioscience), anti-mouse CD103 (2E7), anti-mouse CD107a (1D4B), anti-mouse CD107b (Mac-3) and anti-mouse TNF (MP6-XT22) (all Biolegend). Live-Dead Fixable Green and Live-Dead Fixable Aqua Dead Cell stains was purchased from Thermofisher (Boston, USA).

The following antibodies were used for coating of culture plates for in vitro T cell activation: anti-human CD3 (clone HIT3*α*; Biolegend), anti-human CD28 (clone CD28.2; Biolegend), anti-mouse CD3 (clone 2C11; BioXCell) and anti-mouse CD28 (clone 37.51; BioXCell).

### Mice

C57BL/6J mice were purchased from The Jackson Laboratory. The Rnf144a knockout mouse line was generated using the CRISPR/Cas9 method^64^. Briefly, two sgRNAs (AGCAAGGTACCGGCCCACCT and GCAGGACACCAGTGGGTCGA) were designed to cut near the N-terminus of the mouse RNF144A protein, and were cloned into an sgRNA vector using OriGene’s gRNA Cloning Service (Rockville, MD). These plasmids were then used as templates to synthesize sgRNAs by in vitro transcription using the MEGA-shortscript T7 Kit (ThermoFisher). Cas9 mRNA was transcribed in vitro using the mMESSAGE mMACHINE T7 Ultra Kit (ThermoFisher) with plasmid MLM3613 (Addgene #42251) as a template. Next, Cas9 mRNA (100 ng/μl) was mixed with the two sgRNAs (50 ng/μl each) and comicroinjected into fertilized eggs collected from B6 mice. The injected zygotes were cultured overnight in M16 medium at 37°C in 5% CO2. The next morning, embryos that had reached 2-cell stage of development were implanted into the oviducts of pseudopregnant foster mothers (Swiss Webster, Taconic Farm). The mice born to the foster mothers were genotyped using PCR and DNA sequencing. Where indicated, mice from het x het breeding pairs were weighed weekly after weaning and body composition (fat mass and lean mass) was measured using the EchoMRI™ (Houston, Tx).

### Cell isolation, culture and treatment

Control and KD cell lines were maintained in RPMI 1640 medium (Invitrogen) and HEK293T cells in DMEM (Invitrogen), supplemented with 10% fetal bovine serum (FBS, Invitrogen), 2mM L-glutamine and penicillin-streptomycin (100 IU/ml and 100μg/ml, both from Sigma-Aldrich, UK). The control and KD cell lines were maintained under antibiotic selection in 2 μg/ml puromycin (Invitrogen). For stimulation of cell lines, 100U/mL IL-2 (Roche Diagnostics or Biolegend) was used throughout. Mouse cells were cultured in RPMI 1640 medium supplemented with 10% fetal bovine serum (FBS, Invitrogen), 2mM L-glutamine and penicillin-streptomycin and 2 mM β-mercaptoethanol (Sigma Aldrich).

Mouse splenocytes were isolated by passage through a 40μm filter followed by ACK (Quality biologicals) lysis of red blood cells. Naïve CD4^+^ T cells from spleens and lymph nodes of 6-8 week old mice were purified by negative selection and magnetic separation (Miltenyi). Where indicated, naïve CD4^+^CD25^−^CD62L^+^CD44^−^ cells were FACS sorted using a FACSAria II (BD). CD8^+^ T cells from spleens of mice were isolated by negative selection and magnetic separation (Miltenyi) followed by FACS sorting of naïve CD8^+^CD62L^+^CD44^−^ cells using a FACSAria II.

Preparation of bronchoalveolar lavage fluid (BALF) cells and lung cells was performed according to a published protocol^65^ with minor modifications. Briefly, BALF cells were collected from the lungs of anesthetized mice by gently washing with 1 ml of ice cold PBS containing 2% FBS with 18G plastic cannulae and 1 ml syringes. For isolation of lung cells, mice were perfused with 20 ml of PBS to remove blood from lungs after collection of BALF cells. Isolated lungs were placed in RPMI-1640 containing 2% (vol/vol) FBS, 50μg/ml Liberace TM (Roche, 05401127001), 10 μg/ml DNase I (Sigma-Aldrich, DN25) (4 ml per mouse), then processed with gentleMACS Dissociators (Miltenyi) using gentleMACS C Tubes (Miltenyi), followed by incubation for 45 min at 37°C. Cell suspensions were filtered through 70μm cell strainers, washed twice then purified with 30% Percoll (GE Healthcare).

For restimulation of cells from influenza-infected mice, 1 × 10^6^ cells were incubated with influenza Gp33 peptide (Genscript) at a final concentration of 05μg/mL in the presence of fluorescence-conjugated antibodies against CD107a and CD107b together with Brefeldin A for 5h before additional flow cytometric staining.

### Influenza infection

Where indicated, mice were intranasally infected with 10^3^ EID_50_ of recombinant influenza A/PR/8/34 (H1N1) expressing the LCMV gp33-41 epitope in NA (PR8-33) ^66^.

### Flow cytometry

All flow cytometry was carried out in a final staining volume of 100μL, with data acquisition on an LSR II, LSRFortessa or FACSVerse (all BD Biosciences) within 24h. Appropriate internal controls, isotype controls and Fluorescence Minus One (FMO) controls were used to assign gates. Rat anti-mouse CD16/CD32 (clone 2.4G2; BD) was used for Fc blockade in mouse flow cytometry experiments. FACS data were analysed using FlowJo (Tree Star Inc., Oregon). For Intracellular staining, BD Cytofix/Cytoperm^™^ plus Fixation/Permeabilization Solution Kit was used according to manufacturer’s instructions. For cytokine staining, 4h re-stimulation with PMA (50ng/mL) and ionomycin (1mM) (both Sigma) in the presence of Brefeldin A was carried out prior to fixation and permeabilization. Foxp3 staining was carried out using the kit from eBiosciences as per manufacturer’s instructions. For phospho-flow experiments, cells rested overnight in complete medium (human cells) or X-Vivo 15^™^ (mouse cells) (Lonza) were activated with IL-2 (100U/ml) for stated time points at 37°C, then fixed with BD Cytofix buffer (BD), permeabilized with BD Phosflow Perm Buffer III (BD) and stained with appropriate antibodies as per manufacturer’s instructions.

### Cytokine measurement in cell culture supernatants

IFN*γ* was measured in supernatants of cell cultures stimulated for 16h with IL-2, using the human T_H_1/T_H_2 Cytometric Bead Arrays (BD) as per manufacturer’s instructions.

### Cytotoxicity assays

Cytotoxicity assays were performed as previously described in the presence of IL-2 ^67^. Specific cytotoxicity was calculated using the formula: % specific release = (experimental – spontaneous release) ×100/(maximum – spontaneous release).

### Cell transfections and Luciferase reporter assay

HEK293T cells were transfected using the calcium phosphate method to reconstitute IL-2R signaling, using mammalian expression plasmids for IL-2R*β, γ*c, JAK3, control pRLTK-luciferase (Promega Corp), STAT5-driven luciferase reporter, pPRRIII-CMV-luciferase, pCISTAT5b ^37^ and pFLAGRNF144A. Luciferase activity was measured using the Dual-Glo luciferase Kit (Promega Corp.), according to manufacturer’s instruction. Data were expressed as relative expression of firefly to renilla luciferase. For RAF ubiquitination experiments, 2ug GFP-RAF plasmid, 1ug HA-ubiquitin and 2ug pFLAG-RNF144AWT or C198A mutant plasmids were transfected. Cells were treated with 20μM In-solution MG132 (Calbiochem) for 2hrs prior to cell lysis, where indicated, followed by immunoprecipitation of cell lysates with anti-GFP antibody. In the IL-2-withdrawl experiments IL-2R component plasmids were transfected as above, along with 0.5 μgs pCiSTAT5a, with or without 2μgs pFLAGRNF144A. 48hrs later cells were stimulated with 50 Units/ml IL-2 for 20 minutes and then the cytokine containing medium was removed and replaced with fresh medium containing 20nM JAK inhibitor I (Calbiochem), or not, for the indicated times, followed by whole cell lysates preparation using SDS-sample buffer. One-tenth of each sample was used for western blot analysis. All transfection experiments were repeated at least three times and a representative experiment shown.

### RNA silencing

For production of stably transduced YT cell lines, vector stocks of vesicular stomatitis glycoprotein (VSV-G) pseudotyped lentiviral vectors (pGIPz) encoding shRNA targeting human RNF144A (KD) and non-specific (Control) (all Open Biosystems)were prepared by calcium phosphate mediated three plasmid transfection of HEK293T cells as previously described ^68^. YT cells (2 × 10^6^ cells) were transduced with KD or control lentiviral supernatant and stable cell lines selected in puromycin. For RNF144A knock-down in primary human CD4^+^ T cells, vector stocks of lentivirus encoding shRNA targeting human RNF144A (KD) and non-specific (Control) were prepared by calcium phosphate mediated three plasmid transfection of HEK293T cells. Virus particles were collected and concentrated by precipitation with PEG-it Virus Precipitation Solution (5x) (System Biosciences) and used to transduce pre-activated primary human CD4^+^ T cells in combination with Transdux (System Biosciences). GFP^+^ cells were FACS sorted 48h later, rested overnight and subjected to 2μg/mL puromycin selection before use in functional experiments.

### Immunoprecipitation and Western blotting

Total Cell lysates were prepared using NP40 lysis buffer (150mM NaCl, 0.5%NP40, 50mM Tris, pH 8.0) containing protease inhibitors (Protease V cocktail, Calbiochem or Halt-protease, Invitrogen), 1mM sodium orthovanadate (Sigma), and for ubiquitination reactions 5mM N-ethyl maleimide (NEM, Calbiochem) was included. Total cell lysates were pre-cleared on protein A or G sepharose (Invitrogen) for 1h at 4°C, then lysates were incubated with specific or control antibody for 1h at 4°C followed by incubation with protein A or G for another hour. Immunoprecipitates were washed 3 times with NP40 lysis buffer and boiled in SDS sample buffer. Total lysate or immunoprecipitated samples were resolved on 10% SDS–PAGE gels and transferred to Immobilon PVDF membrane (Millipore). Western blots were performed using primary antibodies directed against proteins of interest, followed by the appropriate species secondary HRP conjugated antibodies, then developed using Clarity MAX™ Western ECL substrate (Bio-RAD) and visualized on Imagequant LAS 4000 instrument (GE). In experiments where specific phosphoprotein followed by total protein detection was performed (eg. pERK/ERK, pSTAT5/STAT5, pRAF/RAF), the blots were probed with phospho-specific primary antibody first and, following ECL, stripped using Restore Plus western blot stripping buffer (Thermo Fisher Scientific) and re-blotted to detect the total specific protein. Quantification of blots was performed using Imagequant software (GE).

### Cloning of Wildtype RNF144A and generation of mutant constructs

The human *RNF144A* cDNA was amplified by PCR from pSPORT6 RNF144A plasmid (Open Biosystems) using primers GCGGGATCCCCATGACCACAGCAAGGTACCGG (P1) and CCGCTCGAGGCGCTTCCTCTAGGTGGGTAACGG (P2), and cloned into the BamHI and XhoI sites of pGex5X3 (Clontech) for N-terminal GST fusion protein generation. For expression into cells with an N-terminal FLAG-tag, *RNF144A* cDNA was amplified using primers CGAGATCTGATGACCACAGCAAGGTACCGG (P3) and CCGGTCGACGCGCTTCCTCTAGGTGGGTAACGG (P4) and cloned into BglII and SalI sites of pFlag-CMV2 (Sigma). FLAG-RNF144A C198A was generated by site-directed mutagenesis using Quickchange (Stratagene) with primers X and X. The FLAG-RNF144A-RING1 (R1) mutant with Cys20 to Arg and Cys23 to Ser mutations in the RING1 domain was made by site-directed mutagenesis using Quickchange (Stratagene) with primers CCGCTGGTGTCTAGAAAGCTCAGTCTTGGGGAGTAC and GTACTCCCCAAGACTGAGCTTTCTAGACACCAGCGG; FLAG-RNF144A-RING2 (R2) with Cys185 to Ala and Cys188 to Ser mutations in the RING2 domains with primers GCGCCCATCAAGCGCGCGCCCAAGAGCAAAGTCTACATCGAG and CTCGATGTAGACTTTGCTCTTGGGCGCGCGCTTGATGGGCGC; the double mutant FLAG-RNF144A-RING1/2 (R1R2) was made by two consecutive rounds of mutagenesis using the R1 and R2 primers. RNF144A^stop249^ was made by PCR using cloning primer P3 together with primer CCGGTCGACGCGCTACTGTGTCCGATGCCAGATCAC to amplify *RNF144A* cDNA with a stop codon at Val250, followed by BglII/SalI cloning into pFlag-CMV2; RNF144A^stop249^ was also cloned into pcDNA3.1 myc/His(-)A to express a protein with a C-terminal myc-tag. This construct was made by PCR using using primers GGAGCGGCCGCCTCAGAGACTGTTCTGCGATGACC and CGCAAGCTTCTGTGTCCGATGCCAGATCAC followed by NotI/HindIII cloning into pcDNA3.1 myc/His(-)A. RNF144A^M140^, which contains an N-terminal deletion and starts at Met140 was made using primers GGAGCGGCCGCACCATGGAATTCTGCT CCACCTGC and cloning primer CGCAAGCTTGGTGGGTAACGGGTCGTCGTC (P6) followed by NotI/HindIII cloning into pcDNA3.1 myc/His(-)A.

### In vitro ubiquitination assays

*E*.*coli* strain BL21DE3(pLysS) (Novagen) was transformed with *RNF144A* constructs cloned into PGEX-5X3 to allow production of GST/RNF144A fusion proteins. Fusion proteins were purified from 5 ml cultures induced with 0.2 mM isopropylthio-*β*-galactoside supplemented with ZnCl_2_ (50 μM) to ensure proper folding of the RING domains. The proteins were purified on glutathione-sepharose beads (Amersham Pharmacia Biotech) after lysis of bacteria by sonication in the presence of 10% Sarkosyl, according to the manufacturer’s instructions. Ubiquitination assays were performed with 10μl glutathione-sepharose beads containing the GST-fusion proteins (∼0.5 μg) in 25 μl reactions containing 100ng E1, 500ng E2, 1μg ubiquitin, 1μg biotinylated ubiquitin, 1mM dithiothreitol [DTT]; 10mM ATP, inorganic pyrophosphatase (IPP, 100U/ml) in reaction buffer (50 mM Tris, pH 7.4; 2.5 mM MgCl_2_;) and incubated for 60min at 30 **°**C, shaking. Ubiquitin, E1 and E2s were purchased from BostonBiochem. The reactions were stopped by adding reducing Laemmli sample buffer and incubating samples for 5min at 95**°**C. Streptavidin-HRP was purchased from Pierce.

### Membrane flotation assays

24h post-transfection, postnuclear supernatants (PNSs) of HEK293T cells were generated by sonicating cells 2 × 15 sec in NTE buffer (20 mM Tris pH 7.4, 150 mM NaCl, 1mM EDTA, protease inhibitors phenylmethylsulfonylfluoride [PMSF, at 1 mM] and CLAP [5 mg/ml each of chymostatin, pepstatin A, antipain hydrochloride and 10 mg/ml leupeptin hemisulphate]) and pelleting nuclei for 10min at 1000 g. PNSs were adjusted to 70% sucrose (w/v) in 2.5 ml, and overlaid with 6 ml of 65% (w/v) sucrose in NTE and 2.5 ml of 10% (w/v) sucrose in NTE. Gradients were centrifuged at 120,000 g for 18 h, following which eleven 1 ml fractions were collected from the top by pipetting. Fraction were adjusted to 1% NP-40, and samples were analyzed by SDS-PAGE and immunoblotting.

### Immunofluorescence

HeLa or Jurkat cells were seeded on 13mm diameter glass coverslips and transfected the next day using Fugene (Roche). Cells were fixed 24 h later with 3% paraformaldehyde, incubated in 50 mM ammonium chloride for 15min, permeabilized with 0.1% Triton TX-100 (Perbio Science) for 5min or not, as indicated, and blocked for 15min in 0.2% gelatin. All solutions were made in PBS. Subsequently, cells were stained with primary and secondary antibodies diluted in 0.2% gelatin for 1h each, followed by three washes in PBS. Coverslips were mounted in Mowiol (Calbiochem-Novachem) and observed by a Leica TCP SP2 AOBSTM confocal microscope. Membrane targeted UD-GFP is GFP fused to the N-terminal unique domain of Lck as previous described ^63^. Lck-GFP is GFP fused to full-length Lck and was a gift from Tony Magee, Imperial College, London.

### Histology and severity scoring

Lung sections were stained with hematoxylin and eosin. Entire slides were digitally scanned using the Nanozoomer Digital Pathology scanner RS 2.0 (Hamamatsu, Bridgewater NJ) and the histology findings evaluated for inflammatory severity. Scores were assigned according to the following scale by an experienced observer blinded to experimental groups: 0, normal; 1, mild changes; 2, moderate changes and 3, severe inflammatory changes.

### RNA Isolation and Quantitative real time PCR

Total RNA was extracted using TRIzol reagent (Invitrogen) and treated with DNAseI (Qiagen). RNA was reverse transcribed to cDNA using iScriptcDNA synthesis kit (Bio-Rad) following the manufacturer’s instructions. Quantitative real-time PCR (qRT-PCR) was performed in triplicate using Taqman® Universal PCR Master Mix (Applied Biosystems) in total reaction volumes of 20 μL and thermocycled in a CFX284 TouchTM Real-Time PCR Detection System (Bio-Rad). Cycle threshold (C_t_) values were exported and normalized against the control probe using the 2^-*Δ*Ct^ method and reported as expression relative to a control condition. The following Taqman gene-specific primer probes were purchased from Applied Biosystems: human *OSM* (Hs00968300_g1), *CISH* (Hs00367082_g1), *LIF* (Hs01055668_m1), *RNF144A* (Hs00207267_m1) and *18S* (Hs99999901_s1), mouse *Rnf144a* (Mm00475644_m1) and *Actb* (Mm00607939_s1). For RNF144A and RNF144B expression in panel S1I qRT-PCR was carried out as previously described ^69^, using the SYBR Green MasterMix (PrimerDesign Ltd, UK) method, in triplicate. Gene-specific primers for RNF144A (F: 5’-GACCTGGCCTGAAGACATTT; R: 5’-TGTGGTCATCGCAGAACAGT) and RNF144B (F: 5’-CCTGACATGGTGTGCCTAAA; R: 5’-CACAGGTACCAAACAGGCAAT) were purchased from Eurofins-MWG, with Human *PPIA* (F: 5’-TGGGCAACATAGTGAGACG; R: 5’-TGTACAGTGGCATGATAATAGC) as reference control.

### RNA-sequencing and microarray

Total CD8^+^ T cells were isolated from the peripheral blood of 2 healthy donors using EasySep™ Human CD8^+^ T Cell Enrichment Kit (StemCell Technologies). Isolated CD8^+^T cells were activated with Dynabeads™ Human T-Activator CD3/CD28 beads (Invitrogen) for 3 days in RPMI medium containing 10% FBS. Cell were then washed and rested overnight in complete RPMI medium without or with JAK kinase inhibitor CP-690550 (Tocris Biosciences), and stimulated for 4 hours with 100U/ml of IL-2. Total RNA from approximately 2 × 10^6^ cells were isolated using the Direct-Zol RNA isolation kit (Zymo Research). RNAseq libraries were prepared as previously described ^70^ using KAPA Standed RNA-seq library preparation kit (KAPA Biosystems), and sequenced on an Hi-Seq 2500 platform (Illumina).

### Computational methods

Gene expression levels was estimated by RSEM software ^71^ as previously described ^72^ from sequenced reads (50bp, single end). The bowtie index for RSEM was created by “rsem-prepare-reference” on all RefSeq genes downloaded from UCSC table browser on April 2017. The differentially expressed genes were identified using edgeR package ^73^. IL-2-regulated genes in CD8^+^ human T cells were defined as those with RPKM >4 with at least 2-fold change at p<0.05 after IL-2 treatment.

Functionally organized GO/pathway term networks from differentially expressed genes were constructed using ClueGO ^74^ for Cytoscape ^75^. A comprehensive list of human E3 ligases and E3 cofactor proteins was obtained from ^76^. RBR family members are from ^77^. A list of deubiquitinase enzymes (DUBs) was obtained form HNGC by using the search term deubiquitinase enzymes and downloading individual datasets for ubiquitin specific peptidases, OUT domain containing, MJD deubiquitinating enzymes, ubiquitin C-terminal hydrolases, MINDY deubiquitinases and merging these with the list of JAMM family members found in Table S1 of ^78^. The combined gene list is shown in **Supplementary Table 4**. Heatmaps of gene expression were constructed using Morpheus (https://software.broadinstitute.org/morpheus/; Broad Institute) throughout. ChIP-seq samples for STAT5B were downloaded from GSM671402 for CD4, and GSM1577746 and GSM1577751 for CD8. H3K27Ac ChIP-seqs were downloaded from GSM1847876 for CD4 and GSM1847878 for CD8. All samples were mapped to human genome, hg19, using bowtie v1.1.2 with -m 1 option. Bound regions (i.e. peaks) were identified by macs14 using default parameters and input controls provided in GSE27158 and GSE64713. Identified peaks were annotated using Homer annotatePeaks. STAT5 bound genes in Figure 1C are defined as those that are shared between two CD8 replicates (GSM1577746, GSM1577751). Tracks were generated using igvtools with the following options [igvtools count -z 5 - w 5 -e 100 --minMapQuality 30] and visualized using IGV browser. Independently called enhancer regions in CD4 and CD8 cells were downloaded from^62^. Gene set enrichment analyses (GSEA) were performed using GSEA version 2.2.2^32^. Gene lists used for GSEA are shown in **Supplementary Table 5**. Transcriptomes of healthy controls and patients with influenza were obtained from GSE101702^38^. ERK pathway genes were sourced from Ingenuity Pathway Analysis (Qiagen). Human IL-2-upregulated genes in **Fig. 1a** (*n* = 1278 genes) were defined, as above, as those with RPKM >4 with at least 2 fold induction at p<0.05 after IL-2 treatment. IL-2-induced and IL-2-repressed genes in **Fig. 2c** and **Supplementary Fig. 4a** were defined as at least 3-fold induced or repressed, respectively, at p<0.05 in wild-type cells on IL-2 treatment. Early *in vivo* CTL effector genes correspond to genes in clusters 1 to 3 from the publication of the ImmGen Consortium^33^.

Estimated probabilities of human gene intolerance to loss of function mutations were obtained from the ExAC database^31^ (n=18,225 genes, release 0.3.1: ftp://ftp.broadinstitute.org/pub/ExAC_release/release0.3.1/functional_gene_constraint/fordist_cleaned_exac_r03_march16_z_pli_rec_null_data.txt; accessed 2016 Aug 18). Multi-species alignment and cladrograms were constructed using CLC Main Workbench 7 (CLCbio, Qiagen).

### Data analysis and visualization

Data were analyzed using Microsoft Excel and GraphPad Prism (Graph Pad Software) and visualized using CLC Main Workbench 7 (CLCbio, Qiagen) and DataGraph 3.2 (Visual Data Tools, Inc). Statistical analyses were performed using appropriate parametric and non-parametric tests as appropriate. Multiple datasets were compared by repeated measures ANOVA. Statistical analysis of data in contingency tables was carried out using the Fisher exact test. A p-value of <0.05 was considered statistically significant throughout.

