## Supplementary Figures 1-9 for "RNF144A shapes the hierarchy of cytokine signaling to provide protective immunity against influenza"

#### Supplementary Figure 1

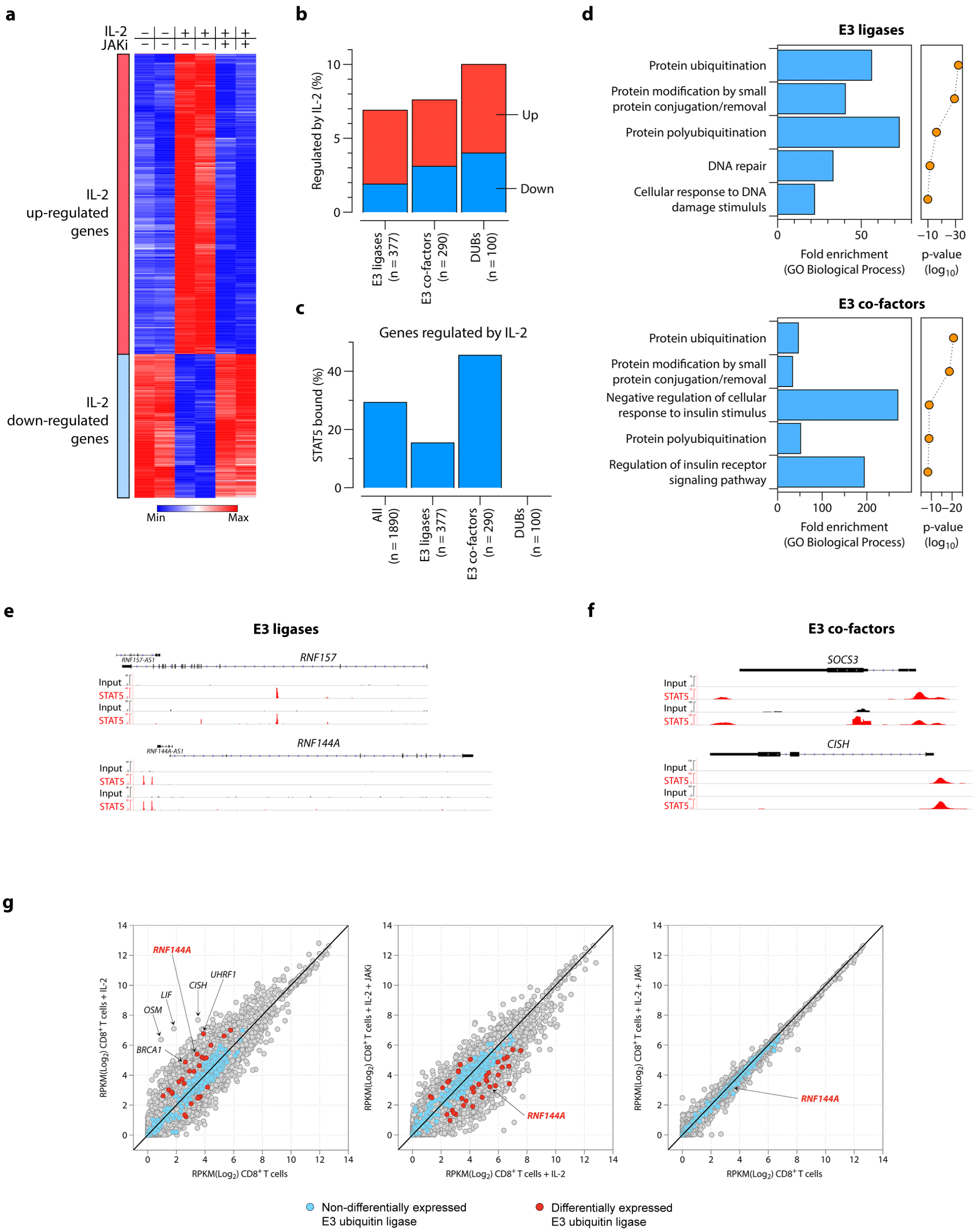

### Supplementary Figure 1

**h**

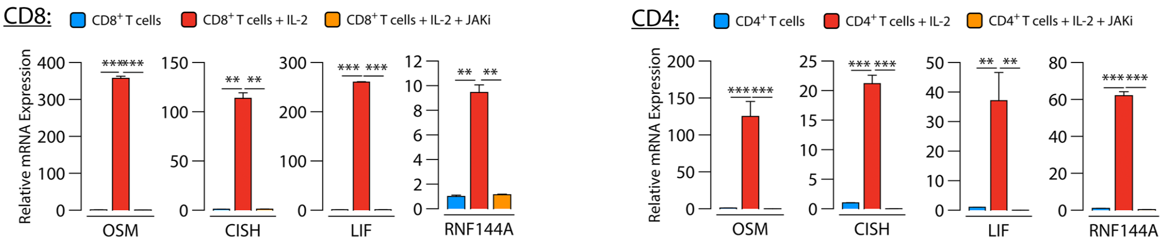


**i**

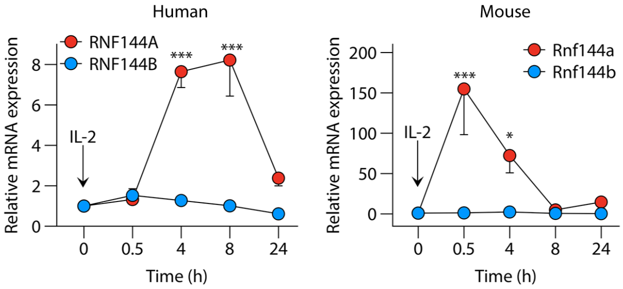


**Supplementary Figure 1. IL-2-induced STAT5 activation regulates *RNF144A*. a-c**, Heatmap of differentially expressed genes in CD8$^+$ T cells (CTLs) activated, or not, with IL-2 in the presence or absence of a JAK inhibitor (JAKi) from *n*=2 donors (**a**), percentage of E3 ligases, E3 co-factor proteins and DUBs regulated by IL-2 (**b**) and the percentage of those bound by STAT5 (**c**). Shown in **b** are IL-2-upregulated genes (red) and IL-2-down-regulated genes (blue). **d**, Top 5 (gene ontology, GO) most significant biological processes associated with E3s and E3 co-factors regulated by IL-2; shown are fold enrichment and p-values. **e-f**, STAT5 ChIP-seq and input tracks for two representative E3 ligases (**e**) and two E3 co-factor protein genes (**f**) bound by STAT5. **g**, Scatterplots of gene expression in CTLs activated, or not, with IL-2 in the presence or absence of a JAKi; each circle represents an individual gene; shown in blue and red are all E3 ligases that are (in red) or are not (in blue) differentially expressed. Shown is one representative experiment from *n*=2 donors. **h**, Quantitative PCR for mRNA of the indicated genes in CD8$^+$ (left) and CD4$^+$ (right) T cells activated, or not, with IL-2 in the presence or absence of a JAKi; shown are pooled data from *n*=3 experiments. **I**, Quantitative PCR time course of *RNF144A* and *RNF144B* mRNA expression induced by IL-2 in human (left panel) and mouse (right panel) CD4$^+$ T cells; pooled data from *n*=3 experiments. Data in (**g-i**) show mean ± sem and have been normalized to baseline (no IL-2 in **h** and time zero in **i**) gene expression levels; *p<0.05 **p<0.01 ***p<0.001.

**Supplementary Figure 2**

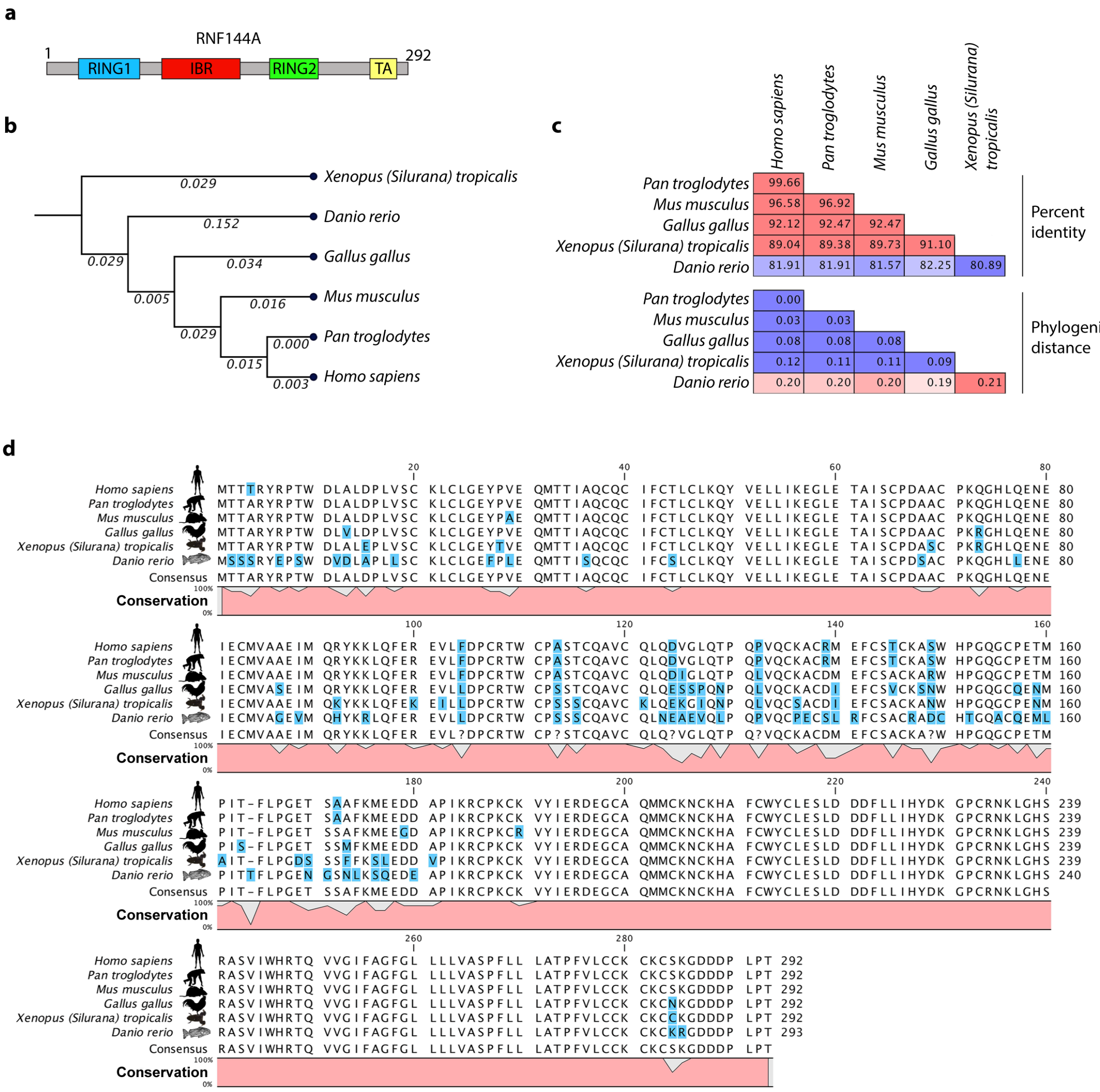


**Supplementary Figure 2. RNF144A is conserved across species. a**, Schematic of RNF144A domain structure; RING, really interesting new gene; IBR, in-between-RING; TA, tail anchor. **b-d**, Phylogenic conservation of RNF144A, showing cladrogram, with branch lengths indicated (**b**), percentage homology (**c**) and amino acid sequence (**d**).

### Supplementary Figure 3

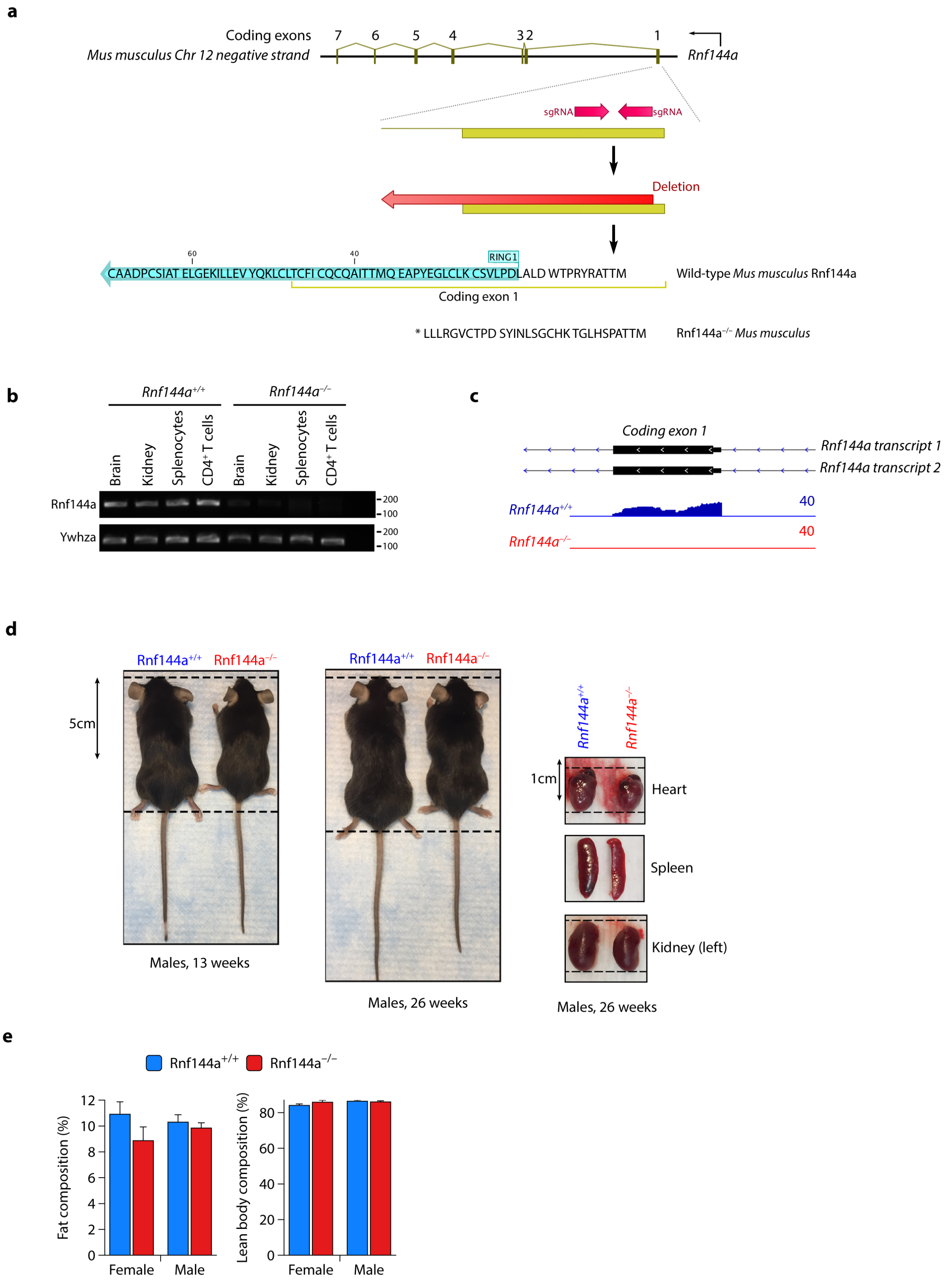

**Supplementary Figure 3**

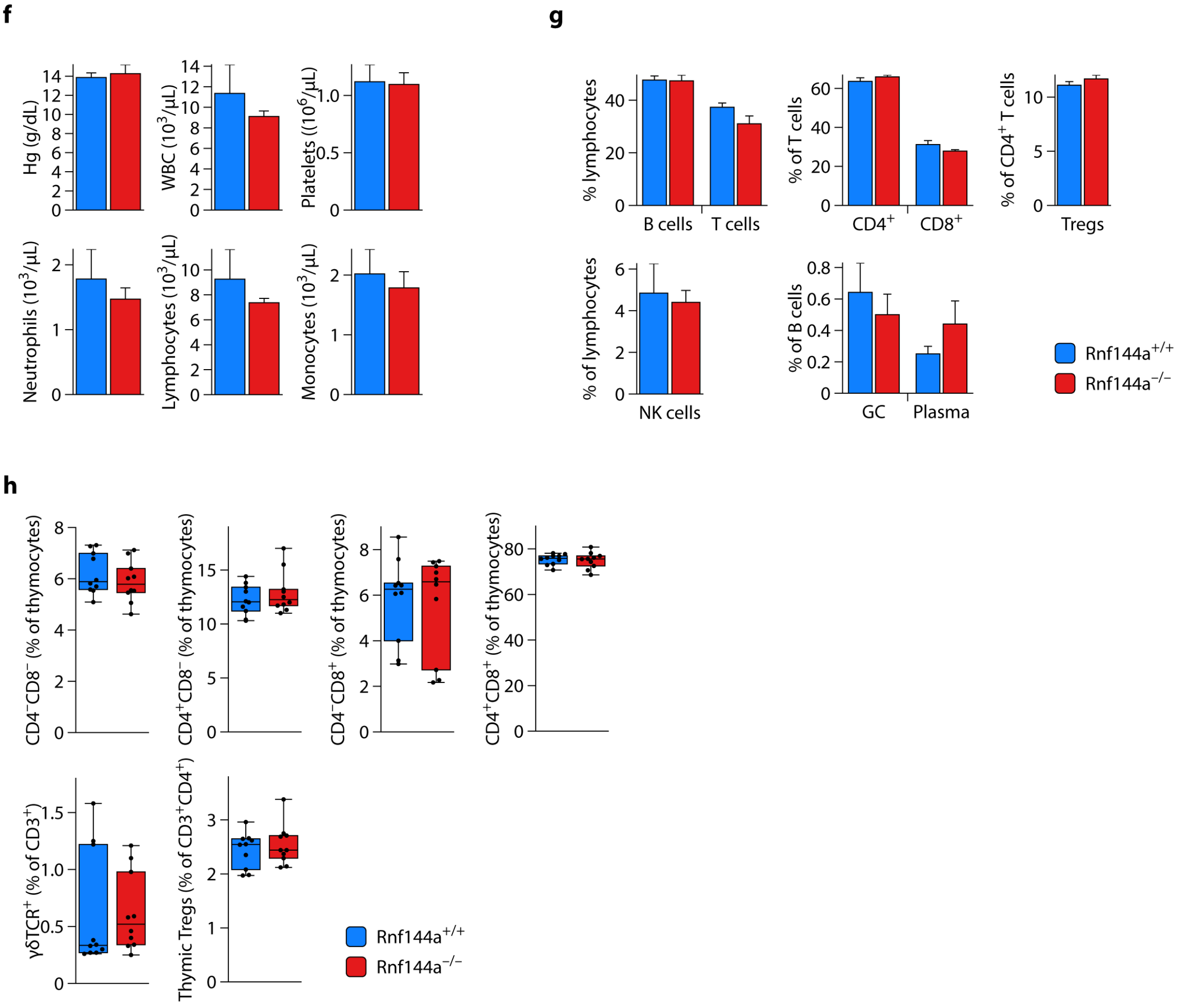


**Supplementary Figure 3. Phenotype of *Rnf144a*$^{-/-}$ mice. a**, Schematic showing generation of *Rnf144a*$^{-/-}$ mice using CRISPR/Cas9; shown are the coding exons of murine *Rnf144a* (top), position of sgRNAs directed against the first coding exon (second from top), resulting genetic deletion in red from the first coding exon to the middle of the next intron (second from bottom) and the wild-type and predicted knockout amino acid sequences (bottom). Marked is the position of the first coding exon in relation to the RING1 domain. **b**, genotyping using primers spanning the first and second coding exons in wild-type and resultant Rnf144a knock-out mice. **c**, RNA-seq tracks from T cells of *Rnf144a*$^{+/+}$ and *Rnf144a*$^{-/-}$ mice showing the first exon. **d**, Representative images of *Rnf144a*$^{+/+}$ and *Rnf144a*$^{-/-}$ mice at 13 (left) and 26 (middle) weeks of age and solid organs harvested at 26 weeks. **e**, Body composition mapping of *Rnf144a*$^{+/+}$ and *Rnf144a*$^{-/-}$ mice showing percentage of fat and lean body mass in both sexes (*n*=5 mice per group per gender). **f**, Complete blood count of *Rnf144a*$^{+/+}$ and *Rnf144a*$^{-/-}$ mice; shown are mean ± sem from *n*=5 mice per group from 2 independent experiments. **g-h**, Immune cell phenotyping of *Rnf144a*$^{+/+}$ and *Rnf144a*$^{-/-}$ mouse splenocytes (**g**) and thymi (**h**); shown are data from *n*=4-7 mice per group from 3 independent experiments. Note that absolute numbers for each cell type were also not different between *Rnf144a*$^{+/+}$ and *Rnf144a*$^{-/-}$ mice.

### Supplementary Figure 4

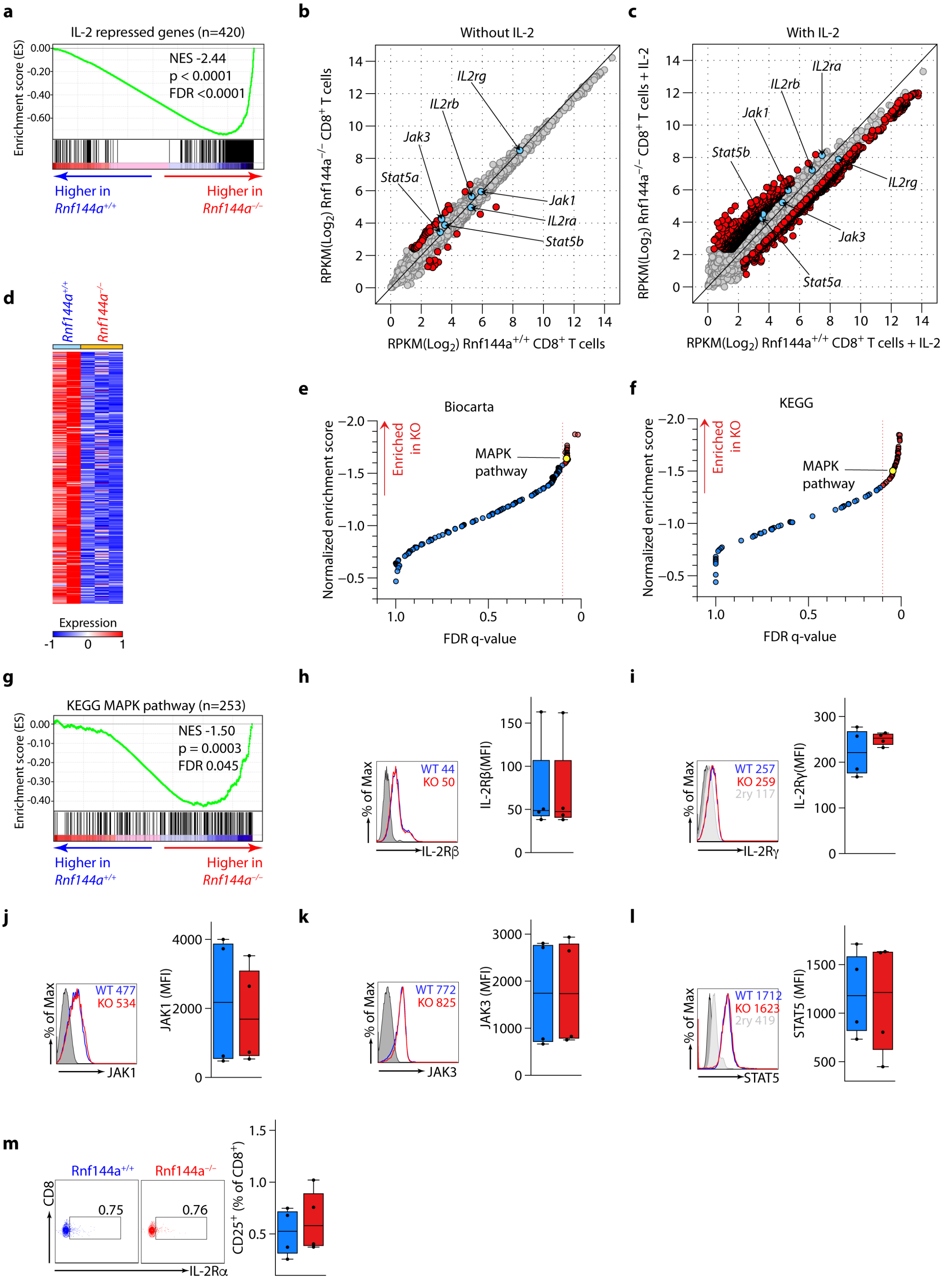


**Supplementary Figure 4. Abnormal IL-2R signal output in *Rnf144a*$^{-/-}$ CTLs.** (**a**) Geneset enrichment analyses (GSEA) of IL-2-repressed (defined as at least 3-fold reduction and p<0.05 in wild-type cells on IL-2 treatment) genes comparing pe-activated IL-2 treated *Rnf144a*$^{+/+}$ and *Rnf144a*$^{-/-}$ CTLs. **b-c**, scatter plots showing gene expression of pe-activated *Rnf144a*$^{+/+}$ and *Rnf144a*$^{-/-}$ CTLs treated (**c**) or not (**b**) with IL-2. Highlighted in red are all differentially expressed genes. Expression of IL-2 receptor components are highlighted in blue, none of which are differentially expressed genes. **d**, Heatmap showing expression of all core enriched (leading edge) genes from **Figs. 2c** and **d**. **e-f**, GSEA output of Biocarta (**e**) and KEGG (**f**) curated pathways comparing IL-2 treated *Rnf144a*$^{+/+}$ and *Rnf144a*$^{-/-}$ CTLs ordered by significance), with pathways enriched in knockout cells at FDR < 0.1 highlighted in red and positions of Biocarta and KEGG mitogen activated pathways highlighted in yellow, respectively. Shown in **g** is the GSEA plot of the KEGG mitogen activated pathway. **h-m**, expression of IL-2Rβ, IL-2Rγ, IL-2Rα, JAK1, JAK3 and STAT5 in *Rnf144a*$^{+/+}$ and *Rnf144a*$^{-/-}$ CD8$^{+}$ T cells. Shown are representative flow cytometry plots (left) and cumulative data (right) from *n*=2 experiments, respectively.

**Supplementary Figure 5**

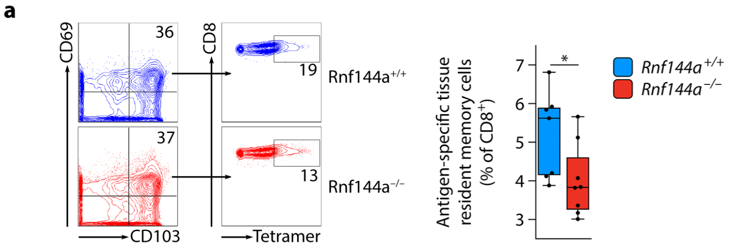


**Supplementary Figure 5. *Rnf144a*$^{-/-}$ mice are immunodeficient. a**, Proportion of antigen-specific tissue resident memory CTLs in lung tissues of *Rnf144a*$^{+/+}$ and *Rnf144a*$^{-/-}$ mice harvested on day 30 after influenza infection; shown are representative flow cytometry plots and cumulative data. *$p<0.05$.

**Supplementary Figure 6**

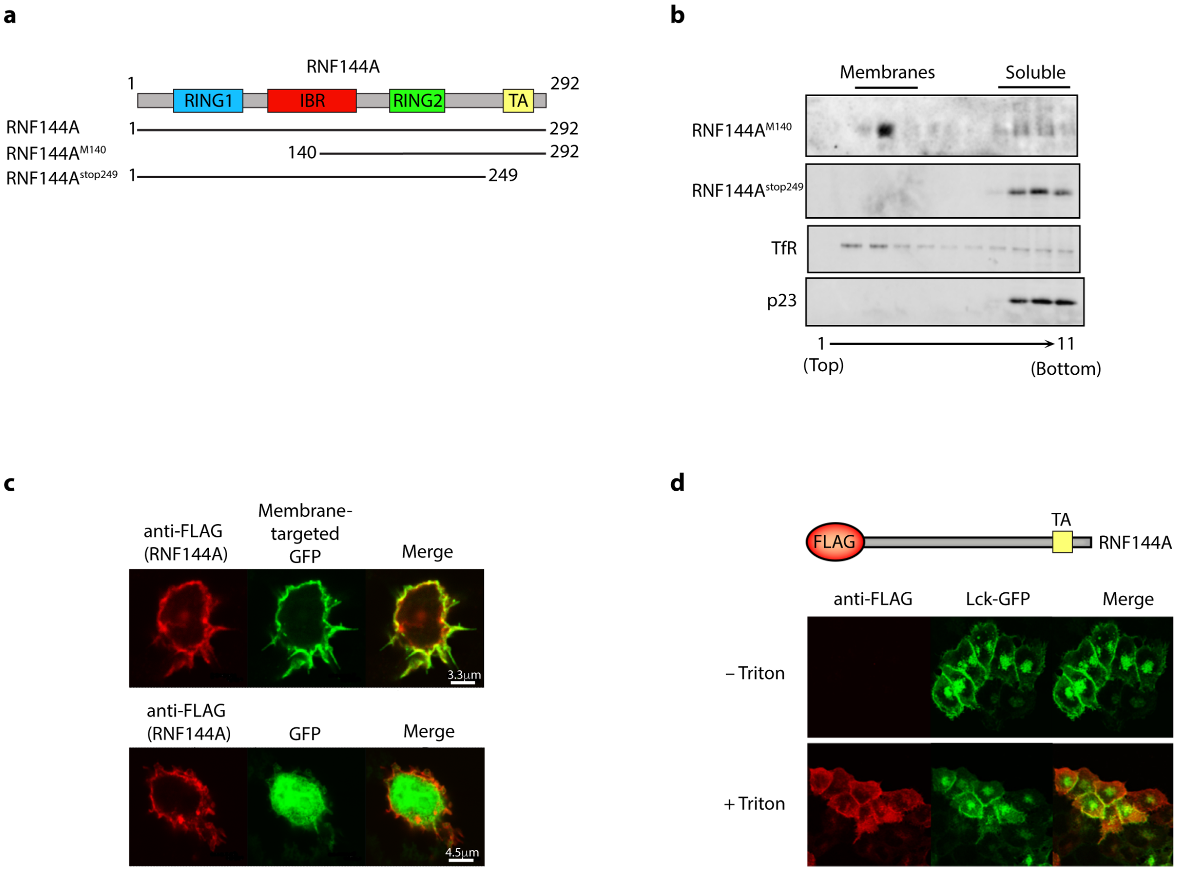


**Supplementary Figure 6. RNF144A is localized to the plasma membrane. a**, Structure of two RNF144A mutant constructs, lacking RING1 (RNF144A$^{M140}$) or tail anchor (RNF144A$^{stop249}$). **b**, Subcellular fractionation on membrane flotation sucrose gradients of HEK293T cells transfected with RNF144A$^{M140}$ or RNF144A$^{stop249}$. **c**, confocal images showing co-localization of FLAG-tagged RNF144A with membrane-targeted UD-GFP[63] (upper panels) but not with nuclear and cytosolic GFP (lower panels) in transfected HeLa cells. **d**, Immunofluorescent staining for FLAG, with (lower) and without (upper) membrane permeabilization in HEK293T cells transfected with N-terminus FLAG-tagged RNF144A and an Lck-GFP fusion protein. Confocal images show FLAG staining (left) overlaid (right) with Lck-GFP (middle). (**b-d**) show representative examples from at least *n*=3 experiments each.

**Supplementary Figure 7**

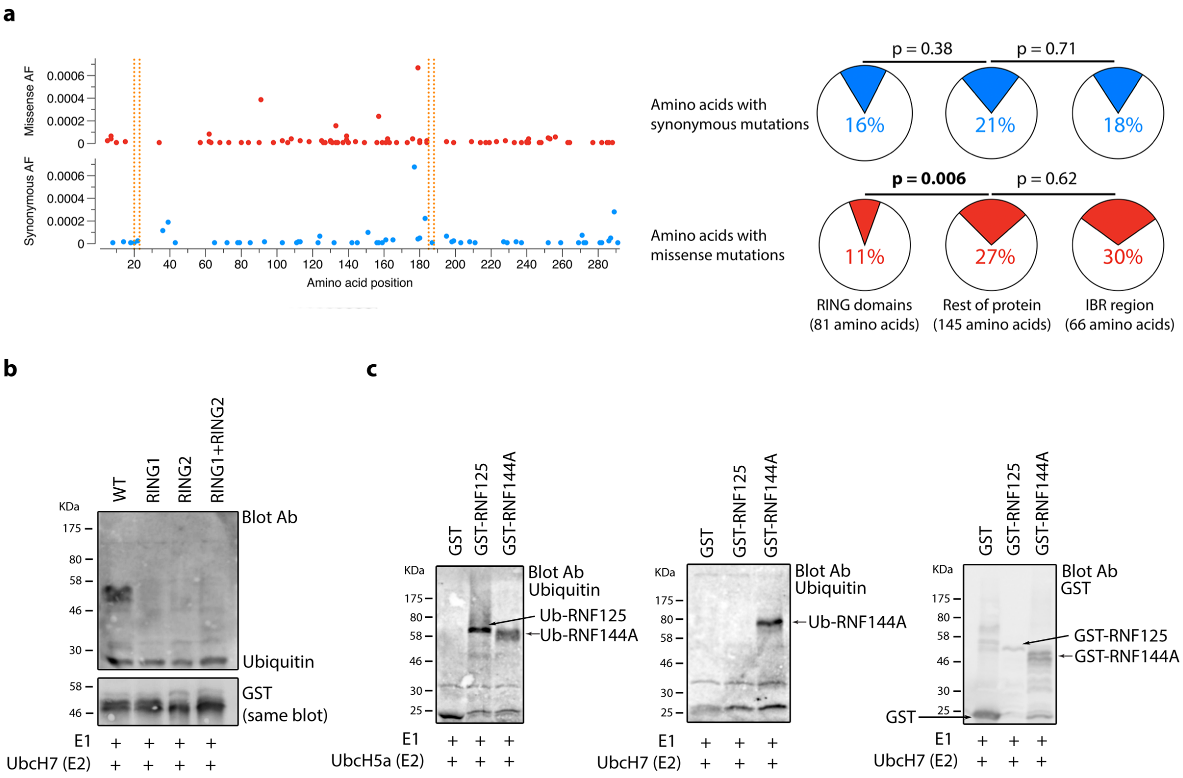


**Supplementary Figure 7. RNF144A has E3 ubiquitin ligase activity.** a, Allele frequency of synonymous (blue) and missense (red) mutations at each residue in RNF144A in the ExAC dataset of whole human exomes (left), with frequency of variants for each region shown on the right; p-values show Fisher exact p-values; dotted orange lines show the position of the conserved Cysteines in RING1 and RING2. The RING domains of RNF144A have unusually low frequencies of missense mutations in human populations and there are no instances of missense variants affecting the conserved cysteines of either RING domain in ExAC, suggesting these to be important structural components of the protein. **b-c**, we expressed and isolated GST-tagged RNF144A in *E. coli* and tested its E3 function *in vitro* in combination with different E2 proteins. **b**, E3 ubiquitin ligase activity with UbcH7 of Wild type (WT), RING1, RING2 and RING1+RING2 mutant RNF144A. **c**, E3 ubiquitin ligase activity of RNF125 and RNF144A with the E2s UbcH5a (left) and UbcH7 (middle); shown on the right is the control anti-GST blot. These experiments show that recombinant wild type GST-RNF144A was able to auto-ubiquitinate *in vitro* with the HECT restricted E2 UbcH7, as well as with the less restrictive E2 UbcH5a, while a control RING E3, RNF125, only functioned with UbcH5a. Thus, RNF144A behaves as a RING-HECT E3 in a manner similar to other RBR family proteins. **b-c** show representative examples from multiple experiments (minimum *n*=3).

**Supplementary Figure 8**

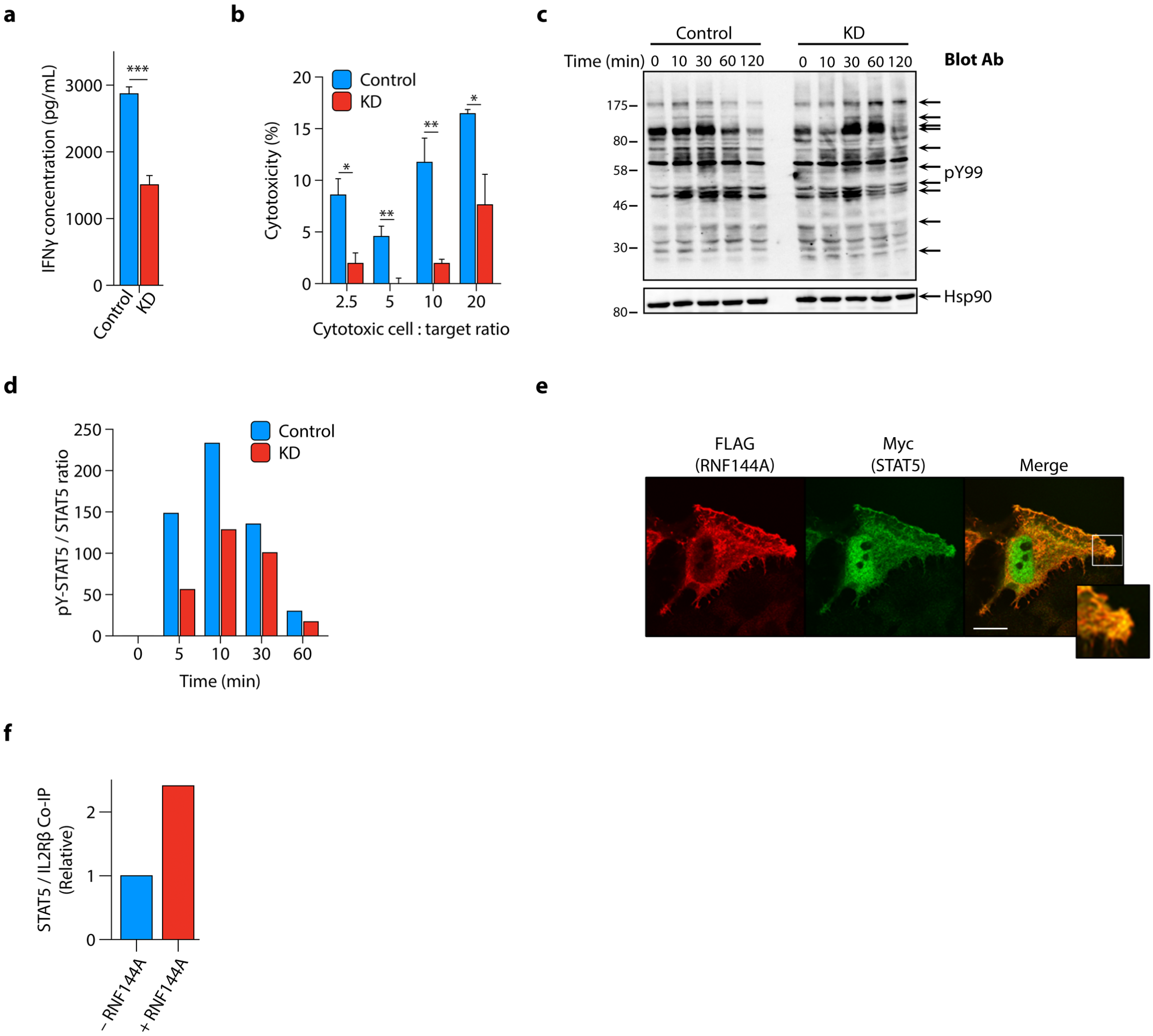


**Supplementary Figure 8. IL-2-induced STAT5 phosphorylation is reduced in the absence of RNF144A. a-b**, IFNγ concentrations in culture supernatants (**a**) and cytotoxicity (**b**) of Control and KD cells stimulated with IL-2. **c**, Global tyrosine phosphorylation (pY99) after stimulation with IL-2 in whole cell extracts of Control and KD cells. **d**, quantification of Western blot in **Fig. 4b**. **e**, Confocal images showing co-localization of FLAG-tagged RNF144A with Myc-tagged STAT5 in HeLa cell; shown are FLAG (RNF144A) staining (left) overlaid (right) with Myc (STAT5, middle); inset highlights overlap between STAT5 and RNF144A at the membrane. The panel shows one representative example from $n$=3 experiments carried out. **f**, Quantification of STAT5 / IL-2Rβ co-immunoprecipitation in **Fig. 4i**. Bars are mean + sem from $n$=6 (**a**) and $n$=4 (**b**) experiments; Western blot in **c** is from $n$=1 experiment. *p<0.05 **p<0.01 ***p<0.001.

**Supplementary Figure 9**

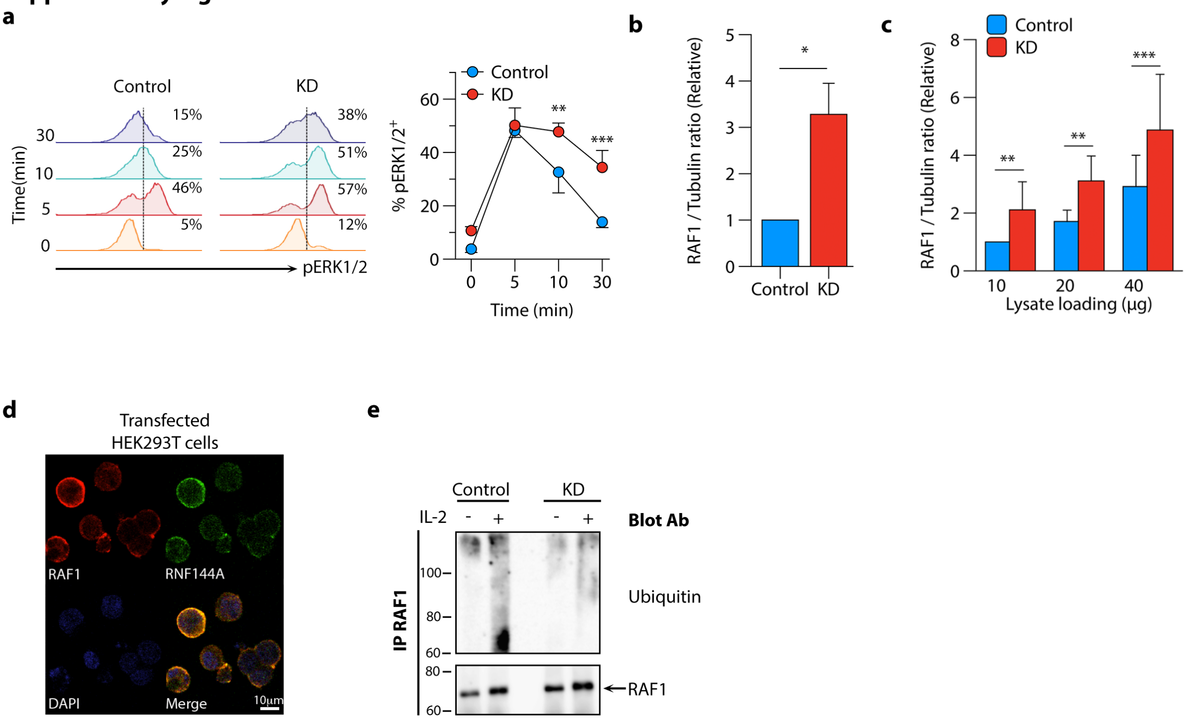


**Supplementary Figure 9. RNF144A restricts ERK MAPK signaling. a**, Flow cytometry histograms of percentage pERK1/2$^{+}$ cells from one representative (left panel) and pooled data (mean ± s.d.) (right panel) from $n$=3 experiments of Control and KD cells treated with IL-2. **b**, Quantification of Western blots of homeostatic (time zero) RAF1 and tubulin in Control and KD cells (**Fig. 5b**) expressed as RAF1:tubulin ratio and normalized to levels in Control cells. **c**, quantification of Western blots of homeostatic RAF1 and tubulin following serial dilution of cell lysates in Control and KD cells (**Fig. 5c**). **d**, Confocal microscopy images of transfected HEK293T cells stained for RAF1, RNF144A and DAPI. **e**, Western blot to detect ubiquitinated RAF1 using anti-ubiquitin antibody (VU-1) from immunoprecipitates of RAF1 from whole cell lysates of Control and KD cells treated with MG132 prior to stimulation, or not, with IL-2. Bars show mean +sem from at least $n$=3 independent experiments throughout. *p<0.05 **p<0.01 ***p<0.001.
